# Rapidly evolving viral motifs target biophysically constrained binding pockets of host proteins

**DOI:** 10.1101/2022.01.29.478279

**Authors:** Gal Shuler, Tzachi Hagai

## Abstract

Evolutionary changes in the host-virus interactome can alter the course of infection, but the biophysical and regulatory constraints that shape interface evolution remain largely unexplored. Here, we focus on viral mimicry of short host-like peptide motifs that allow binding to host domains and modulation of cellular pathways. We observe that motifs from unrelated viruses preferentially target conserved, widely expressed and highly connected host proteins, enriched with regulatory and essential functions. The interface residues within these host domains are more conserved and bind a larger number of cellular proteins than similar motif-binding domains that are not known to interact with viruses.

In stark contrast, rapidly evolving viral-binding human proteins form few interactions with other cellular proteins, display high tissue specificity and their interface residues have few inter-residue contacts. Our results distinguish between highly conserved and rapidly evolving host-virus interfaces, and show how regulatory, functional and biophysical factors limit host capacity to evolve, allowing for efficient viral subversion of host machineries.

## Introduction

As obligatory parasites, viruses must engage in numerous interactions with intracellular host proteins. Due to their small genomes encoding relatively few proteins, viral proteins are thought to be multifunctional^1–3^, but the molecular mechanisms behind this are poorly understood. One mechanism that may support multifunctionality is viral mimicry of host-like peptide motifs (also called **ELMs** – eukaryotic linear motifs). ELMs can interact with diverse host proteins allowing for efficient subversion of host pathways. They function as short interacting modules with peptide recognition domains (**PRDs**^4^), including SH3, WW and PDZ domains, as subcellular localization signals, and as sites for cleavage or for posttranslational modifications^5^ (**Fig 1A**). ELMs are often embedded in intrinsically disordered regions of proteins^6^ – segments that do not adopt a well-defined tertiary structure but that are important for various regulatory functions^7–10^.

**Figure 1:**
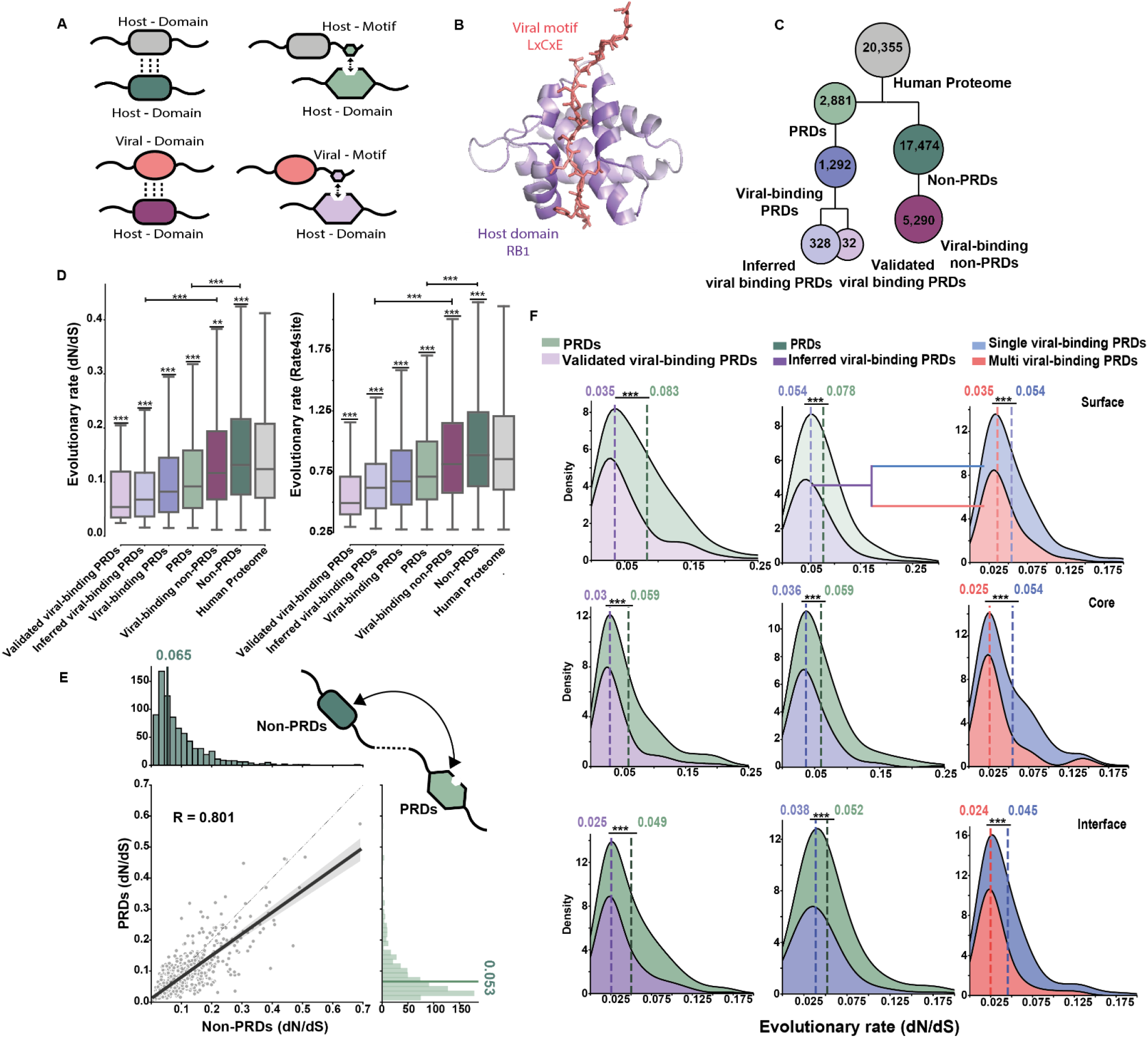
Viral host-like motifs target conserved human proteins. (**A**) A schema of host-host (top) and host-virus (bottom) protein-protein interactions (PPIs) by type: domain-domain (left) *versus* domain-motif (right) interactions. (**B**) An example of a viral motif-host domain interaction. The LxCxE motif of the Simian virus 40 Large T-antigen protein (red) interacts with the host Retinoblastoma-associated protein B domain (purple) (PDB :1GH6). (**C**) Classification of 20,355 human proteins, based on whether they have a PRD (peptide recognition domain) and whether they bind viral proteins. In the set of viral-binding PRDs (1,292 proteins), we inferred that the host-virus interaction is mediated through domain-motif interactions if a matching motif was found in disordered regions of the interacting viral protein (328 proteins). 32 cases of host-virus PPIs are experimentally-validated to be mediated through domain-motif interactions. (**D**) Distribution of evolutionary rates for the sets of proteins mentioned in C. The evolutionary rate is relative across all proteins and is computed based on substitutions across a set of 12 one-to-one orthologs in vertebrates, using either: the ratio between the number of synonymous and non-synonymous substitutions (dN/dS values, left, 7,435 proteins in total) or substitutions per residue-specific rate (using rate4site, 7,445 proteins in total, right). Differences between the distributions were compared using Mann-Whitney test and corrected by FDR. (**E**) Paired distributions of evolutionary rates of domains within proteins, where the Y-axis represents the average value of the PRD and the X-axis represents the average value of the non-PRDs from the same protein (n=807 proteins). PRDs have significantly lower mean rate in comparison to non-PRDs (P-value =1.57×10^−6^, Mann-Whitney test). (**F**) Kernel density plots showing the distribution of evolutionary rates of amino acids at the core, surface and interface of PRDs (left - experimentally validated viral-binding PRD, middle and right - inferred viral-binding PRDs). PRDs are split into those that are not known to be bound by viral motifs (left and middle, in green) those that are known to be bound viruses through domain-motif interaction (left and middle, in purple) and those by those that are known to be bound by a single (right, in blue) and multiple viral motifs (right, in red). Dashed lines and values represent the medians of the distribution. The differences between the distributions were compared using Mann-Whitney test. (*** - P < 0.001, ** -P < 0.01, * P < 0.05). A similar plot of all PRD using the rate4site approach is shown in Fig Supp 5.

These short and simple motifs can readily evolve in viral proteins to facilitate the formation of interactions with host proteins through domain-motif interactions mimicking within-host interactions and displacing them^11–13^. Literature-based surveys^14,15^ and proteome-wide analyses^16,17^ suggested that ELMs are found in a diverse set of viruses and can participate in many stages of viral replication (see for example **Fig 1B**).

Despite their simplicity, ELMs can form complex regulatory modules that enable efficient perturbation of cellular networks^16,18^. For example, a PDZ-binding motif at the terminus of the E6 oncoprotein in cancer-causing papillomaviruses enables interaction with PDZ domains of several human signaling proteins, supporting cell transformation^19^. Furthermore, ELMs have been associated with virulence and adaptation to new hosts, as in the case of glycosylation of influenza hemagglutinin^20–22^.

While previous studies characterized the occurrence, evolution and function of viral motifs, we here focus on the host side: By comprehensively analyzing a large dataset of host-virus domain-motif interactions, we show that viral motifs interact with conserved, widely expressed and highly connected human proteins, enriched with essential and regulatory functions. Importantly, those human proteins targeted by viral motifs are even more conserved and more highly connected than respective proteins with the same domain composition that are not known to bind viruses.

We then contrast these host domain–viral motif interactions with the remaining set of host-virus interactions, and with those interactions where the host proteins display signatures of positive selection. We observe significant differences in evolutionary, regulatory, biophysical and functional characteristics between the groups. Based on this, we suggest a general model of the evolution of the host-virus interactome that accounts for both highly conserved and rapidly evolving interfaces between host and viral proteins.

## Results

### Distinguishing different types of host-virus interactions by domain and motif mapping

We first partitioned the human proteome into two main classes (see **Methods** and **Figs 1A, C**): Proteins that are known to have a domain that binds motifs (these proteins will be denoted as **PRD proteins** hereafter, and their motif-binding domain itself will be denoted as **PRD**) and all other proteins, that are not known to contain such domains (denoted as **non-PRDs**). In each of these classes, we also identified the subset of human proteins with at least one experimentally verified interaction with a viral protein (**viral-binding PRDs** and **viral-binding non-PRDs**). In viral-binding PRDs we inferred the subset of host-virus protein-protein interactions (PPIs) that may be mediated by PRD-motif interactions, by searching for a match between a motif in the viral protein and a PRD in the human interacting protein (**inferred viral-binding PRDs**, see **Supp Fig 1** and **Supp Table 1** for details). We also identified a smaller subset of host-virus PPIs where the motif’s role in the interaction was experimentally validated (**experimentally validated viral-binding PRDs**).

### Viral motifs mediate interactions with conserved human proteins

We next studied the relative evolutionary rates of the above-described group of proteins using two approaches: (1) We calculated dN/dS – the ratio between nonsynonymous and synonymous substitutions – a measure indicative of the selective pressure on coding sequences, using Selecton^23^. This was done for each human gene with a multi-sequence alignment with orthologs from 11 additional vertebrates (see **Methods**). (2) We calculated the relative evolutionary rate of each residue and the average rate of each protein using the same set genes and orthologs as in (1), using Rate4Site^24^ – a method that takes into account the tree phylogeny and the amino acid substitutions observed across the multi-sequence alignment (see **Methods**). Throughout the analysis we observed similar trends using the two approaches and we note that the two measures are significantly correlated despite being based on different metrices (R=0.9, P-value<10^−308^, see **Supp Fig 3**).

We observed that viral-binding proteins tend to be more conserved than non-viral-binding proteins in the same class (this is true for both PRD or non-PRD proteins) (**Fig 1D**). Importantly, viral-binding PRD proteins constitute a highly conserved set: more conserved than both viral-binding non-PRD proteins as well as the entire PRD protein set (FDR-corrected P-value = 4.12×10^−15^ and 8.94×10^−44^, respectively, Mann-Whitney test). Finally, the small set of experimentally validated viral-binding PRDs is the most conserved set of proteins in our dataset (FDR-corrected P-value = 4.31×10^−5^, Mann-Whitney test).

### Binding pockets in human PRDs targeted by viral motifs represent highly conserved regions

We next partitioned the sequences of PRD proteins into segments of different domains, using PFAM domain annotations^25^. For each PRD protein, we contrasted the evolutionary rates of the two types of domains: PRDs *versus* non-PRD domains. We observed that PRDs are more conserved than their respective non-PRDs, suggesting that PRDs evolve slower than other domains in the same protein (P-value = 1.57×10^−6^, Mann-Whitney test, **Fig 1E**). This is true both in cases where the human protein is known to interact with viruses, as well as in cases where such an interaction was not reported (**Fig 1E** and **Supp Fig 4**).

Next, we divided the residues within each PRD to residues predicted to be at the core, surface and interface – the latter are the residues within the domain’s binding pocket that interact with the motif (see **Methods**). Interestingly, when comparing PRDs that are known to form interactions with viral motifs *versus* PRDs that are not known to form such an interaction, we observed that all three regions – surface, core and interface – are more conserved when a viral motif is known to interact with the domain (**Fig 1F** and **Supp Figs 5-6**). This trend is observed both in the smaller set of experimentally validated cases, where the role of viral motifs in mediating interactions was validated, as well as in the larger set of inferred domain-motif interactions (**Fig 1F** and **Supp Figs 5-6**). Additionally, these observations hold when we test each type of PRD independently (e.g., when comparing viral-binding SH3 domains with non-viral-binding SH3 domains, **Supp Fig 7**). In contrast to our expectations, interface residues in PRDs do not display higher evolutionary rates than core residues (showing similar or even lower evolutionary rates, depending on the set compared – see **Supp Fig 6**). This suggests that similar evolutionary constraints are imposed on interface and core residues in the case of PRD proteins.

Finally, we compared core, surface and interface residues between PRDs for which only a single interaction with a viral protein is known, with PRDs that are known to interact with motifs from multiple viruses (**Fig 1F right**, and **Supp Figs 5-6**). Intriguingly, we observed that PRDs that interact with multiple viruses evolve more slowly than the respective PRDs that interact with only one virus. This is again observed in all three structural regions: surface, core, and interface (FDR-corrected P-value = 3.66×10^−5^, 4.36×10^−5^ and 1.37×10^−7^ for surface, core and interface comparisons, respectively, Mann-Whitney test). Thus, proteins from unrelated viral families convergently evolved motifs that interact with extremely conserved binding pockets, even with respect to similar binding pockets of other PRD proteins.

### SARS-Cov-2-specific motif targets a highly conserved Discoidin domain in the human Neuropilin-1 protein

The emerging picture from the above analysis is that host-like motifs that can rapidly evolve in viral proteins^11,14^ interact with constrained binding pockets of host PRD proteins. The asymmetrical nature of this type of host-virus interactions can be seen in a recent finding regarding SARS-Cov-2 interactions with human proteins: SARS-Cov-2 Spike protein has a polybasic furin cleavage site that is absent in other coronaviruses of the same clade^26^. This is important for enhanced SARS-Cov-2 infection and may increase its transmissibility^27^. This cleavage, in turn, exposes a short motif that enables the viral S1 protein to interact with the human Neuropilin-1 protein (NRP1), which increases virus uptake into the cell^28,29^. Here, we analyzed NRP1 protein conservation and observed that it belongs to the third most conserved set of proteins in the human proteome (dN/dS=0.88 *versus* an average value of 0.155 across the entire human proteome; 33.8% of the human proteins have lower dN/dS values than NRP1). Furthermore, we observed that NRP1 residues that interact with the viral motif show complete conservation in all vertebrates used in our analysis, as well as in fruit and insectivore bats (bats are suspected to be the origin of SARS-Cov-2) (**Fig 2**). This analysis exemplifies how fast-evolving motifs can be utilized to interact with a highly conserved host machinery, important for intracellular replication. In the following sections we analyze biophysical factors that likely contribute to the observed evolutionary conservation of PRDs and their binding pockets.

**Figure 2:**
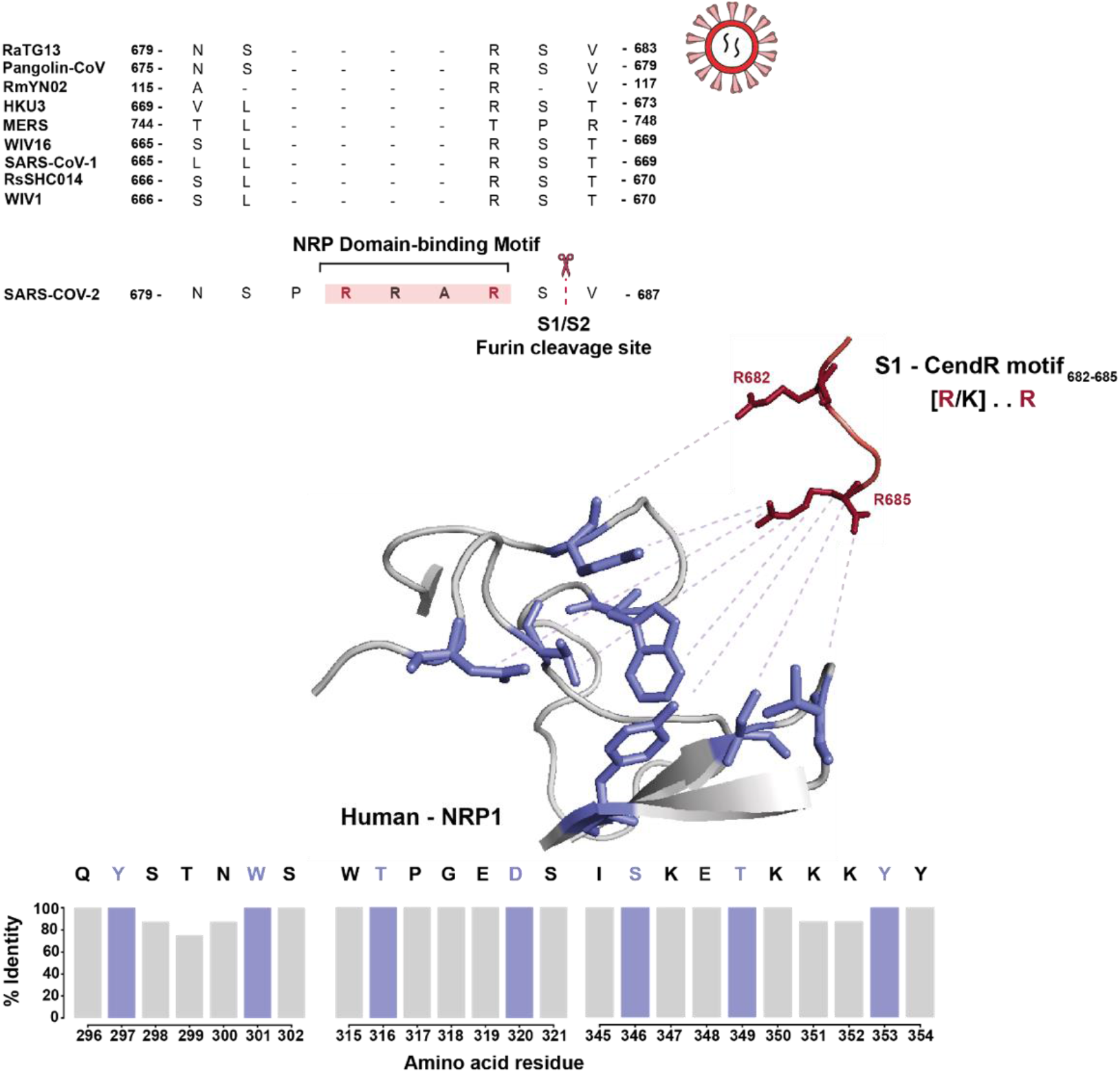
SARS-Cov-2-specific motif binds to the highly conserved binding pocket with Neuropilin Receptor-1 Domain. **Top**: MSA based on Spike glycoprotein of 10 selected Coronaviruses. In red - CendR motif and adjacent S1/S2 furin cleavage site of SARS-Cov-2. **Middle**: 3D structure of Neuropilin-1 (NRP1) domain – SARS-Cov-2 CendR motif interaction. The [R/K]..R motif of SARS-Cov-2 (in red) interacts with Human NRP1 (in purple and ribbons). Dashed lines represent the interactions between the residues (PDB: 7JJA). **Bottom**: percentage of identity of residues in the binding pocket of NRP1 across vertebrates, based on an MSA of 8 orthologs from ENSEMBL genomes used in this study and two bat species (Human, Chimpanzee, Mouse, Pig, Horseshoe bat, Megabat, Chicken and Tortoise). In purple – residues in NRP1 domain that interact with the SARS-Cov-2 motif. See comparison with respective distribution of the percent identity in Supp Fig 8.

### Viral motifs target interface regions that are binding platforms for numerous human proteins

Following this, we asked whether additional biophysical, cellular and functional characteristics distinguish viral-binding PRD proteins from other proteins. We first compared the number of within-host PPIs (interactions formed between two human proteins) – between the group of viral-binding PRDs and other human proteins, using the curated set of the human interactome from STRING^17^. We observed a clear trend where PRD proteins have a significantly higher number of PPIs within the human proteome (FDR-corrected P-value = 3.31×10^−164^, **Fig 3A**), in agreement with reports on SH3- and PDZ-containing proteins^4,30^. Importantly, the set of viral-binding PRDs has a particularly high number of PPIs – significantly higher than both the general set of PRD proteins as well as the set of viral-binding non-PRDs (FDR-corrected P-value = 7.69×10^−9^ and 6.75×10^−50^, respectively, Mann-Whitney test).

**Figure 3:**
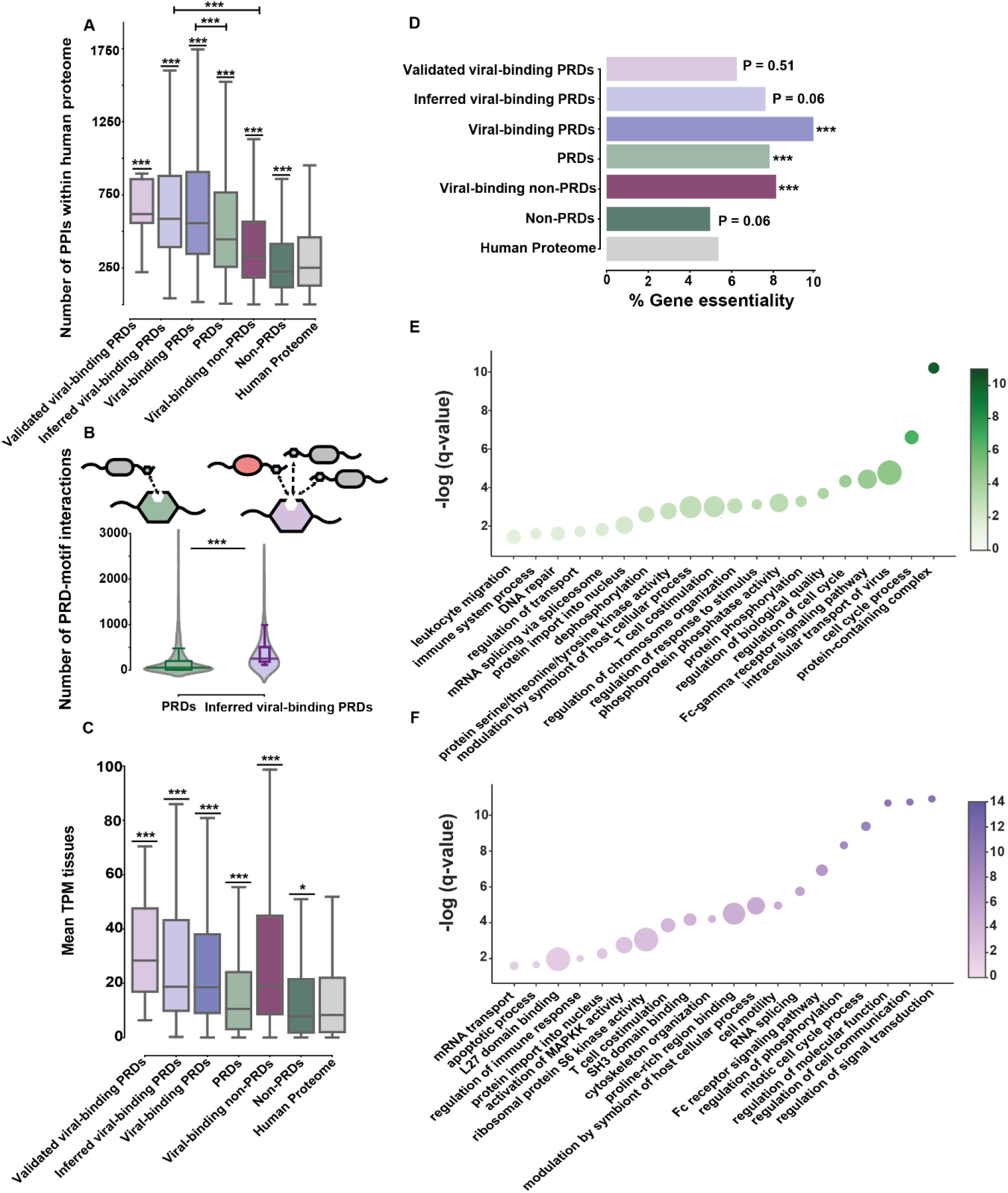
Viral host-like motifs target widely expressed and highly connected essential human proteins. (**A**) Distribution of the number of within-host PPIs for the sets of proteins defined in 1C (n = 13,999 proteins with PPIs from STRING^70^). Differences between the distributions were compared using Mann-Whitney test and corrected by FDR. (**B**) A comparison of “binding pocket usage” between all PRD proteins (n = 2,881, green) and the subset of RBDs known to bind viral motifs (n = 328, purple). The differences between the distributions were compared using Mann-Whitney test. (**C**) Distributions of average gene expression level across healthy adult human tissues (n = 19,670). Differences between the distributions were compared using Mann-Whitney test and corrected by FDR. (**D**) The percentage of essential genes out of each group of proteins defined in 1C. Enrichment with respect to the whole-human proteome was computed using Fisher’s exact test and corrected by FDR. (**E**) Bubble plots of functionally enriched GO terms in viral-binding PRDs in comparison with the rest of the PRD protein set. The size of each bubble is determined by enrichment level and the color gradient by –(log(q-value)). (**F**) Bubble plots of functionally enriched GO terms in viral-binding PRDs in comparison with viral binding non-PRDs. (*** - P < 0.001, ** -P < 0.01, * P < 0.05).

We next calculated how many motifs embedded in different human proteins interact with each domain. For this, we searched for a matching co-occurrence of a PRD and its cognate binding motif in two human proteins that are known to interact (e.g., by searching for a match between an SH3 domain and an SH3-binding motif in two human proteins that are known to physically interact, see **Methods** and **Supp Fig 1**). This gives a measure of “binding pocket usage” – indicating how many human proteins interact with the PRD using the same motif and forming an interaction with the same binding pocket.

We then divided the set of PRD proteins into those that are known to bind viral motifs and the rest of the PRD proteins. When we compared the “binding pocket usage” measure of the two PRD protein classes, we observed that viral-binding PRDs have significantly higher number of binding motif occurrence than the non-viral-binding PRD proteins (P-value = 3.2×10^−26^, Mann-Whitney test, **Fig 3B**). Thus, viral motifs preferentially interact with those pockets of PRDs that are heavily used as an interaction module by a multitude of human proteins.

In summary, PRD proteins that viruses interact with, serve as important hubs within the cellular network, and their motif-binding pockets form numerous interactions with other human proteins. In this manner, viral motifs bind evolutionarily constrained pockets within PRDs that serve as interfaces central to the human interactome and can potentially disrupt the interactions of a large number of cellular proteins.

### Viral motifs target highly expressed essential human proteins

Next, we studied another factor known to be associated with conservation – gene expression level, by calculating the average transcript expression level across human tissues using the human protein atlas database^18^. We observed that PRD proteins in general, and their subset that binds viral proteins in particular, tend to be more highly expressed across tissues in comparison with other groups of proteins (FDR-corrected P-value = 4.22×10^−10^ and 2.92×10^− 91^, respectively, Mann-Whitney test, **Fig 3C**).

We next focused on gene essentiality measures using three different screens for essential genes^31^ (see **Methods**). We observed that a significantly higher fraction of viral-binding PRDs are essential genes (9.98% in the set of viral-binding PRD proteins), in comparison with the entire human proteome (5.37%, FDR-corrected P-value = 6.254×10^−10^, Fisher’s exact test). The set of viral-binding PRD proteins is enriched with essential genes also in comparison with the set of all PRD proteins and the set of viral-binding non-PRDs (9.98% versus 7.84% and 8.12%, P=value = 0.014 and 0.023, Fisher’s exact test, respectively) (**Fig 3D**).

Thus, the set of viral-binding PRD proteins is associated with functional, biophysical and regulatory constraints that are reflected in high conservation of the protein sequence in general, and the binding pocket of the PRD in particular (**Fig 1D-F**).

### Viral motifs target regulatory proteins associated with assisting or inhibiting viral replication

We next searched for functional enrichment of viral-binding PRDs by using Gorilla, a tool that enables gene enrichment analysis by contrasting two different groups of proteins^32^. We first asked which functions are enriched in the set of viral-binding PRDs in comparison with the rest of the PRD proteins (**Fig 3E**). We observed that viral-binding PRDs are enriched with regulatory functions that control the immune system, including “leukocyte migration” and “regulation of response to stimulus”. In addition, viral-binding PRDs are enriched for processes that are known to be modulated by viruses, including “cell cycle”, “DNA repair” and “intracellular transport of viruses”. Thus, in comparison to the general set of PRDs, those PRDs that are known to be targets of viral proteins tend to be enriched with regulatory immune functions or with pathways that are either inhibited or hijacked by the virus to assist in their replication.

When we compared the set of viral-binding PRD proteins with viral-binding non-PRDs, we observed an enrichment in various regulatory pathways, including signal transduction, phosphorylation, regulation of immune response and regulation of cell communication (**Fig 3F**). In addition, viral-binding PRDs are enriched with particular pathways that are relevant to viral life cycle and its modulation of cellular homeostasis, including “cell motility”, “cytoskeleton organization”, “apoptotic process” and various mRNA-related processes.

In summary, the set of proteins to which viruses bind to through domain-motif interactions is involved both in regulation of processes that assist in virus replication, as well as in regulation of antiviral defense. This is observed both in comparison to non-PRD proteins that are bound by viruses as well as in comparison to other PRD proteins that are not known to interact with viruses (see the complete list of GO terms **in Supp Tables 2-3**).

### Viral-interacting PRD proteins are depleted of signatures of positive selection

Following our analysis that showed that viral-binding PRDs and their interfaces are evolutionarily conserved, we asked what characterizes viral-binding proteins that are rapidly evolving: We focused on genes that exhibit signatures of positive and turned to delineate the cases in which they interact with viral proteins. Previous work showed that various antiviral proteins as well as host factors – host proteins that virus subvert for their replication – display signatures of positive selection, and that this rapid coding sequence divergence of the interacting host protein can impact virus ability to infect different host species^33^. We used a previous analysis of 11,152 human proteins, of which 330 proteins had strong statistical evidence for positive selection based on a comparative analysis of their orthologs across 9 primates^34^ (**Fig 4A**). We observed that proteins with signatures of positive selection comprise 2.96% of the general set of the human proteome. However, only 1.57% of the PRD proteins and 0.88% of the inferred viral-binding PRDs have evidence for positive selection – a significant depletion (FDR-corrected P-values of 0.001 and 0.046, respectively, Fisher’s exact test). Furthermore, none of the experimentally validated viral-binding PRDs have any evidence for positive selection.

**Figure 4:**
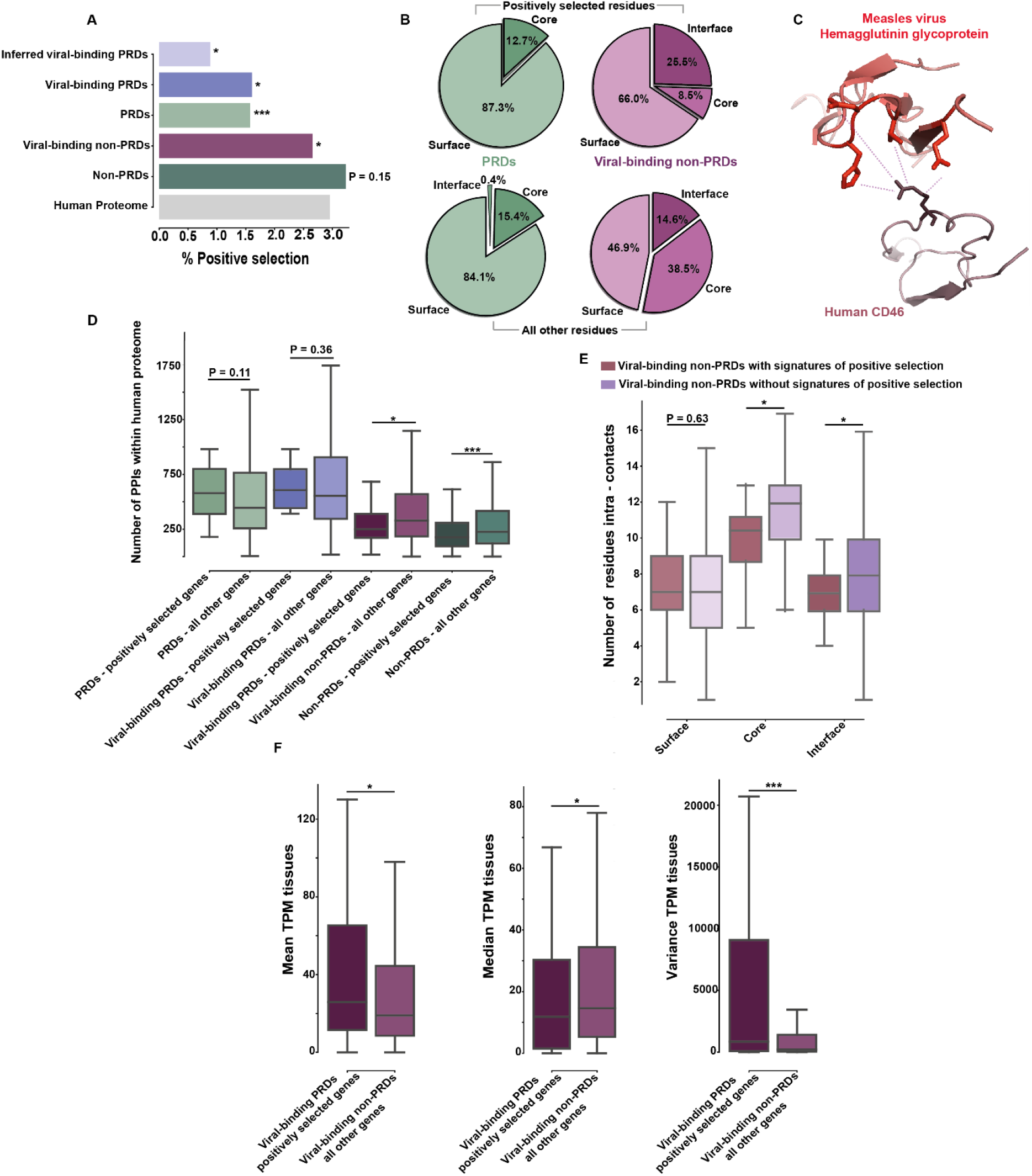
Positively selected viral binding proteins are depleted of PRDs, have low connectivity and are tissue-specific. (**A**) The percentage of proteins that display signatures of positive selection out of each group of proteins defined in 1C. Enrichment with respect to the whole-human proteome was computed using Fisher’s exact test and corrected by FDR. (**B**) Top - Pie charts showing percentage of positively selected residues in surface, core and interface of (1) PRD proteins (left, green, n=110 residues in 29 PRD proteins, none of them is located at the PRD interface), and (2) viral-binding non-PRDs (right, purple, n=141 residues in 36 proteins with PDB structures). Bottom - Pie charts showing percentage of all other residues in surface, core and interface of (1) PRD proteins (left) and (2) viral-binding non-PRDs (purple). (**C**) An example of host-virus interaction with positive selection: Human CD46 (purple) interacting with Hemagglutinin glycoprotein of the *Measles virus*, with the positively selected Arg69 and its viral interacting residues (in ribbons). Dashed lines represent the interactions between the residues (PDB: 3INB). (**D**) Distribution of the number of within-host PPIs for the sets of proteins defined in 1C, partitioned into those with and without signatures of positive selection. Differences between the distributions were compared using Mann-Whitney test and corrected by FDR. (**E**) Distributions of interactions formed between residues within the same protein (intra-contacts) in surface, core and interface; between viral-binding non-PRDs, with and without signatures of positive selection (red and purple, respectively). The differences between the distributions were compared using Mann-Whitney test and corrected by FDR. (**F**) Comparisons of measures of cross-tissue gene expression between viral binding non-PRDs with and without signatures of positive selection: Left - mean TPM values across the tissues. Middle - median TPM values across the tissues. Right – variance of TPM values. The differences between the distributions were compared using Mann-Whitney test and corrected by FDR. Positively selected genes show lower median and higher variance in comparison with other genes, see additional analyses in Supp Fig 12 (*** - P < 0.001, ** -P < 0.01, * P < 0.05).

### Unlike highly conserved PRD-motif interfaces, viral-interacting non-PRD protein interfaces are enriched with positively selected residues

Next, we asked whether interfaces between PRDs and viral motifs contain residues that are positively selected, by exploring the spatial location of positively selected residues within the structures of 29 PRD proteins that have signatures of positive selection (5 of these interact with viruses through domain-motif interactions). None of the 110 positively selected residues in PRD proteins are predicted to be part of the interface between the domain’s binding pocket and the motif’s interacting residues: 96 of these residues are predicted to be at the surface and the remaining 14 are found at the core (**Fig 4B left, in green**).

In stark contrast to interfaces between host PRDs and viral motifs that are devoid of rapidly evolving residues, interfaces between viral proteins and host non-PRDs are enriched with positively selected residues: 25.5% of the positively selected residues within human non-PRDs are predicted to be at the interface with viral proteins – a significant enrichment with respect to the fraction of all interface residues in these proteins – 14.7% (P-value = 0.004, Fisher’s exact test, **Fig 4B right, in purple**).

In summary, PRD proteins, including viral-binding PRD proteins, are depleted of signatures of positive selection. The few positively selected residues in PRD proteins are not located at the interface with motifs. An opposite trend is observed in non-PRD proteins where their interfaces with viral proteins are enriched with such rapidly evolving residues.

### Positively selected non-PRD proteins that bind viruses are depleted of interactions with other human proteins

We next focused on the number of interactions formed between proteins with signatures of positive selection and other human proteins: We observed that viral-binding non-PRD proteins with signatures of positive (there are 88 such cases, see for example **Fig 4C**) form significantly fewer interactions with other human proteins in comparison to all other non-PRD proteins (FDR-corrected P-value = 0.012, Mann-Whitney test, **Fig 4D** and **Supp Fig 10**). Interestingly, we did not observe such a trend with positively selected PRD proteins (either with or without known interactions with viral proteins). This result may stem from the fact that in PRD proteins many of the PPIs are mediated by domain-motif interactions, whereas the positively selected residues of these proteins are not found in the binding pockets that mediate these interactions.

### Rapidly evolving residues within viral-binding non-PRD proteins have few interactions with other residues

Next, we focused on residue-level analysis and studied the interactions formed between residues within the same protein. We used proteins with solved structures and calculated the number of non-covalent interactions formed between residues (“**intra-contacts**”) using the CSU method^15^. We compared the numbers of intra-contacts that are formed by residues with and without signatures of positive selection in viral-binding non-PRD proteins: We observed that positively selected residues are significantly depleted of intra-contacts with respect to other residues both at the core and at the interface (FDR-corrected P-value=0.045 for both core and interface intra-contacts, Mann-Whitney test, **Fig 4E**). Surface residues do not show significant differences in their intra-contact numbers between these two classes of residues, in line with the relatively flexible nature of surface residues and the few residues they usually interact with. We extended this analysis to proteins without experimentally solved structures, by using the structures predicted by AlphaFold^35^. We observed the same trend in core and surface of predicted structures as well, although this was not statistically significant (**Supp Fig 11A**). This suggests that rapidly evolving regions within viral-binding human proteins have relatively few contacts with other residues within their domain, which likely supports the rapid rate of amino acid substitutions observed in these positions across species.

Interestingly, when looking at binding pocket residues of PRDs – the residues that form the interface with motifs, we observed that the number of contacts formed between them and other residues within the PRD domain is higher than the corresponding number of residue contacts at the interfaces of non-PRD proteins (FDR-corrected P-value = 1.46×10^−24^, Mann-Whitney test, **Supp Fig 11B**). Furthermore, when considering the interactions formed between the residues in the binding pocket and the residues in the motif (“**inter-contacts**”), the combined total number of interactions that residues in the PRD binding pockets have is even higher than the number of contacts formed within the core of these PRDs (FDR-corrected P-value = 7.77×10^−61^, Mann-Whitney test, **Supp Fig 11B**). This high connectivity of residues in the binding pocket of PRDs is in line with their slow evolutionary rate reported above (**Fig 1F**). In summary, rapidly evolving residues found at the interface between non-PRD proteins and viral proteins, have few contacts with other residues, in contrast to interface residues of PRD proteins that are highly connected to other residues.

### Positively selected viral-binding human proteins are expressed in a tissue-specific manner

We next studied the expression profiles of viral-binding non-PRDs that have signatures of positive selection in comparison with other viral-binding non-PRDs, using cross-tissue gene expression data^36^. Interestingly, we observed that viral-binding non-PRDs with signatures of positive selection are highly expressed, but only in a subset of tissues. This can be observed by comparing the mean, median and variance distributions of expression of viral-binding non-PRDs, with and without signatures of positive selection (**Fig 4F** and **Supp Fig 12**). This suggests that those rapidly evolving proteins that form interactions with viral proteins, tend to be tissue-specific and likely have specific functions that are important in those tissues, rather than across a wide spectrum of tissues. Thus, fewer regulatory constraints are likely imposed on viral-binding non-PRDs with signatures of positive selection as manifested in their narrow expression profile.

### Positively selected viral-binding human proteins belong to two separate functional classes

Finally, we examined what functions are associated with these 88 viral-binding non-PRD proteins that have signatures of positive selection: Using GO term enrichment analysis with the g:Profiler program^37^, we observed that nearly ∼36% of the genes (32 genes) have antiviral or immune-related functions. These include, among others, several **restriction factors** – proteins that act to inhibit viral replication by directly binding viral proteins or other viral elements, such as tetherin (BST2) that efficiently prevents the release of diverse enveloped viruses by tethering their virion to the cellular membrane^38,39^.

The remaining set of 56 genes (∼64% of 88 genes) has no enriched functions and likely represents a diverse group of proteins that during viral infection act as **host factors** – a group of unrelated proteins that are modulated by viruses to support their replication. This set includes, for example, the cytosolic chaperon DNAJB14, that assists in mobilization of nonenveloped viruses, such as simian virus 40, out of the ER^40,41^ (see **Supp Table 4** for the complete list of 88 PSGs that are viral-binding non-PRDs and their functional classifications).

Our analysis of viral-binding proteins that have signatures of positive selection suggests that this set can be divided into two distinct functional classes: host factors and host restriction factors, that play opposite roles during viral infection (supporting and inhibiting viral replication, respectively). However, in both cases, these proteins interact with relatively few other host proteins. Moreover, their positively selected residues form few contacts with other residues, alleviating constraints associated with connectivity. In contrast, such constraints are observed in the case of the highly conserved PRD interfaces that mediate binding through domain-motif interactions.

## Discussion

Previous studies showed that residues found at the interface between host and viral proteins often evolve rapidly as a result of their antagonistic interactions^34,42,43^. In contrast, other studies observed that viral proteins preferentially bind conserved human proteins^44–47^. Our analysis suggests a more complex landscape of host-virus interactions and elucidates several principles in the evolution of these interfaces.

Biophysical and regulatory factors, including the number of interacting proteins, the structural network of contacts between residues, gene essentiality, expression level and functional pleiotropy, shape the evolution of viral-binding proteins and their interface residues. These factors can limit host capacity to evolve in response to deleterious interactions with viral proteins, conferring an advantage to viral proteins in hijacking vital host machineries. Viral mimicry of short host-like motifs represents an extreme of such cases, where the introduction of a few mutations in the viral proteins leads to formation of these motifs, enabling them to interact with highly conserved and highly connected residues in important regulatory proteins. In addition, PRDs serve as binding modules for numerous host proteins, enabling viruses to perturb numerous cellular interactions using few viral motifs. For example, a recent study on SARS-Cov-2 nucleocapsid protein showed that its interaction with the human G3BP protein through a short motif leads to displacement of several stress granule-resident proteins, effectively disrupting stress granule formation^18^. Notably, our analysis shows that those PRDs that viral motifs are known to interact with are more conserved and interact with larger numbers of cellular proteins in comparison with PRDs that have the same domain but are not known to interact with viruses. This suggests a general trend where motifs from unrelated viruses preferentially interact with highly-connected PRD proteins.

When comparing viral-interacting PRDs with viral-binding proteins with signatures of positive selection, we repeatedly observe contrasting trends. Proteins from the latter group interact with a small number of human proteins and their interface residues have few inter-residue contacts. Furthermore, these rapidly evolving viral-binding proteins tend to be expressed in a tissue-specific manner and to be enriched with functions unrelated to regulation and signaling that are the hallmarks of viral-binding PRDs.

This comparative analysis enables us to disentangle host-virus interactions and to suggest a model for their evolution (**Fig 5**): Viruses engage in numerous interactions with intracellular host proteins. Some of these human proteins are restriction factors that function as inhibitors of the viral lifecycle in various ways, while others are host factors – proteins that are hijacked by the viral machinery to act as accessories for viral replication. Both types of interactions represent a genetic conflict where the interaction confers an advantage to one side but is unfavorable to the opposing side. To avoid this interaction, mutations can emerge in the disadvantaged partner. From the host perspective however, these escape mutations cannot be fixed in cases where they are deleterious to the host protein function, such as when they abrogate its interactions within the human interactome.

**Figure 5:**
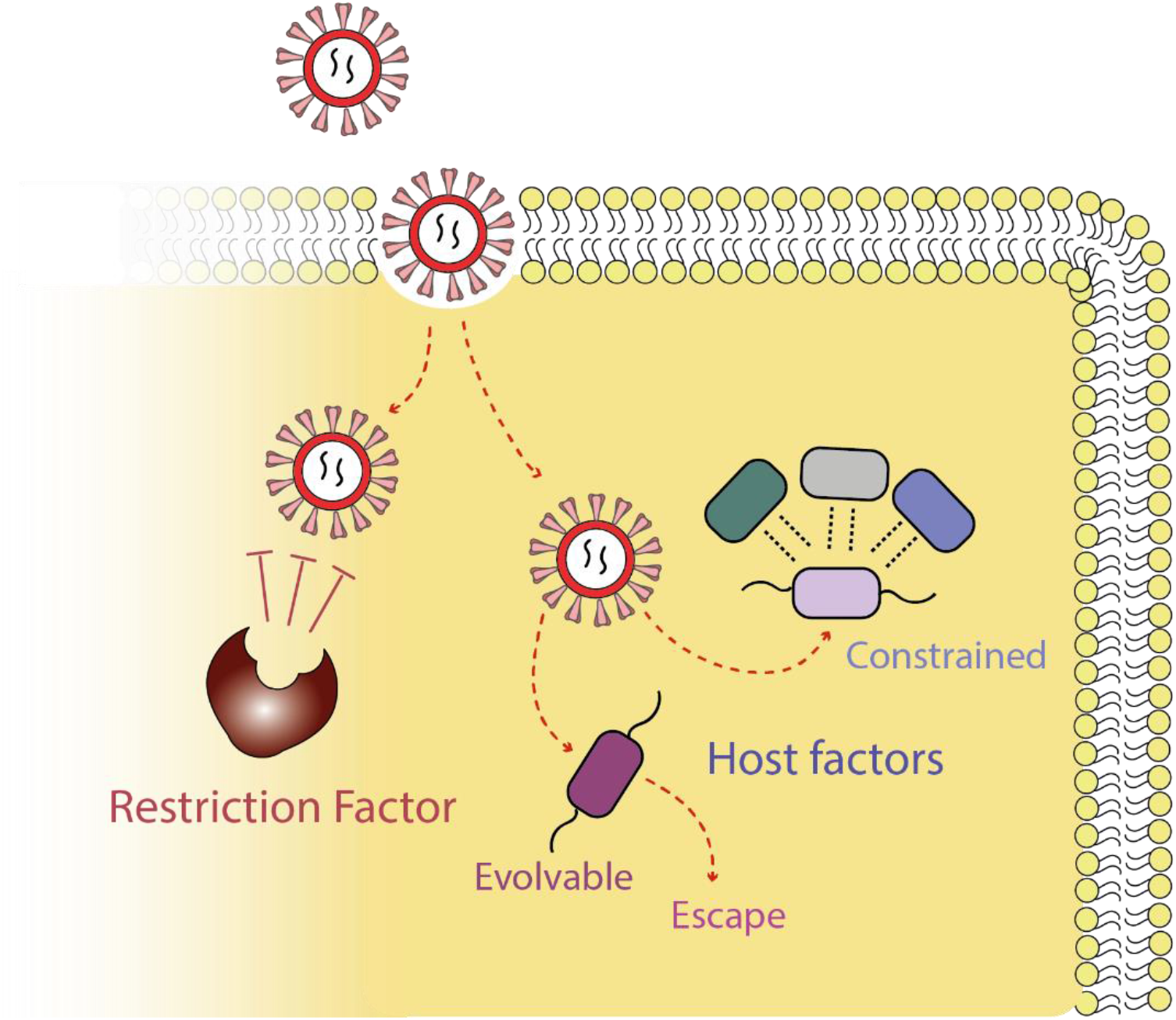
Proposed model for the evolution of the host-viral interactome. Viral proteins form intracellular interactions with host proteins that are either restriction factors (left, in brown) or host factors (right), that inhibit or support viral replication, respectively. The interfaces formed between host factors and viral proteins may be under strong selective pressure due to various constraints (such as in the case of PRD binding pockets). Alternatively, they can evolve rapidly in cases where their residues do not constitute an interface with many human proteins or in cases where these proteins are under weaker regulatory or functional constraints (center). Host factor escape from interactions with viral proteins results in an overall depletion of rapidly evolving proteins in the set of human-viral interactome.

Host-virus interactions involving restriction factors that often function exclusively in viral inhibition, display patterns of rapid evolution (**Fig 5, left**). The genes encoding restriction factors exhibit signatures of positive selection as these proteins predominantly function in the antiviral response, are usually not involved in other cellular pathways, and are often upregulated in response to infection.

The case of host factors that are subverted by viruses to assist in their replication, and that represent a larger fraction of host-virus interactions, is more complex: Some of these host factors may evolve rapidly and these cases are mostly attributed to proteins that do not form many interactions with other host proteins. In these cases, their residues are relatively free to evolve (**Fig 5, centre**). At least in some cases, rapidly evolving genes are exclusively expressed in a few tissues, in contrast to highly conserved proteins that are widely expressed across tissues (**Figs 3C *vs*. 4F**). This tissue-specific expression pattern, observed in the set of positively-selected viral-binding non-PRDs, further suggests that there are fewer functional constraints on these genes, supporting their rapid evolution.

In other cases, however, host factors have stronger constraints since they function in pathways unrelated to immunity, are widely expressed, or interact with various human proteins in the same regions that are also bound by viral proteins (**Fig 5, right**). In these cases, the viral protein engages in interactions with host proteins that cannot easily escape from these interactions. PRD proteins exemplify this scenario at its extreme – in these cases, viral proteins bind the host protein using motif mimicry in the same manner and in the same region that is used for binding by numerous host proteins^14,17^. These PRD binding pockets are thus highly constrained since escape mutations that prevent viral-binding can compromise important interactions of the PRD with its interacting human proteins. Since, as we observed here, PRD proteins are highly expressed in a large number of tissues and are enriched with essential functions, these mutations may have a strong deleterious effect on host fitness.

An outcome of the above-described dynamics can be that over long evolutionary timescales, there will be an enrichment for interactions of viral proteins with conserved host factors, which we indeed observed (**Fig 1D**). In the course of host evolution and as part of the selection imposed by viral invasion, some of the less evolutionarily constrained host factors will escape viral interactions (**Fig 5, centre**). In contrast, constrained host factors will not be able to escape such interactions, leading to their enrichment in the host-viral interactome. Since we usually characterize host-viral PPIs in a single evolutionary “time point” (that of human) – some of the evolvable host factors still interact with viral proteins while other evolvable proteins escape them, depending on the state of the evolutionary arms race between the host and the virus in these particular PPIs. The escape of some of the evolvable host factors, leads to their depletion in the set of host-virus interactome.

Our study reveals a complex evolutionary landscape of the interactions between viruses and their hosts, where regulatory, functional and biophysical constraints shape the evolution at the interface between the two antagonistic partners. Our results distinguish between highly conserved and rapidly evolving interfaces, raising new questions about the impact these interactions have on virus ability to transfer to new host species, as well as to how this evolution affects host response to contain infection from emerging viruses.

## Methods

### Dataset assembly and protein classification

As the basis of our analyses, we used the non-redundant set of human coding genes from Ensembl version 102^48^. For each gene, we took the longest coding transcript. We linked ENSEMBL ID with UniProt ID^49^ using ENSEMBL BioMart tool, to allow searching for domains in these proteins using Pfam database that is UniProt-based (**see Supp Table 1**). We used the Pfam-A list^25^ (last modified on 1/5/2020) to extract the domain composition of each of the human proteins.

Proteins were defined as containing peptide recognition domains (**PRD proteins**) if they have at least one domain that is defined as a PRD, or an ELM binding domain, based on a list of 142 such domains that was downloaded from the ELM database^50^ (downloaded on 22/03/2021). All other human proteins were defined as **non-PRD proteins**.

To find human proteins that are experimentally validated to physically interact with viral proteins, we used the **HVIDB database**^51^ that includes 48,643 experimentally verified human-virus interactions assembled from six different databases (PHISTO^52^, PDB^53^, HPIDB^54^, VirusMentha^55^, IntAct^56^ and VirHostNet^57^) and additional curations. We discarded interactions with missing protein information in UniProt or that are not curated as meditated by physical association, association or direct interaction between the human and the viral proteins, leaving 48,027 host-virus PPIs. Proteins that have at least one known interaction with a viral protein were defined as **viral-binding PRDs** if they have at least one PRD and **viral-binding non-PRDs** otherwise.

The list of viral-binding PRDs may contain human proteins that bind viral proteins independently of their PRD. We thus searched within the set of viral-binding PRDs those human proteins that we can infer that they are likely to interact with the viral protein through domain-motif interactions. Since each type of PRD has one or more motifs that are known to specifically bind to them^50^, we searched in the interacting viral protein sequences for segments that match the motif (e.g., we searcher for co-occurrence of an SH3 domain and an SH3-binding motif in two interacting proteins). **Inferred viral-binding PRDs** were defined as PRD human proteins where a match was found between their PRD and a matching motif sequence in the interacting viral protein. To search for motif sequences, we used the motif’s regular expression as defined by the ELM database. We filtered motif-matching segments that were not found in disordered regions. For this, we used the IUPred program^58^ (version 2A, with default parameters and with the ‘long’ option), and defined the segment that matches the motif as disordered if its residues had an average IUpred score of 0.4 or higher. The result of this filtration was a set of 5,064 unique host-virus PPIs that are inferred to be mediated by a viral-mimicking motif, embedded within a disordered region, and a host PRD (see **Supp Fig 1** for a scheme of this workflow). We note that in some cases functional ELMs may be found in relatively structured surface regions^59,60^, such as in the case of glycosylation sites. However, we here choose to use a strict threshold and analyze a smaller subset of inferred motifs to have fewer false positives.

Finally, we collected a set of 52 **experimentally validated viral-binding PRDs** – a set of host-virus PPIs that the mechanism of interaction through domain-motif binding was experimentally verified (for example, by point mutation). This set was assembled from the curated ELM database^50^ and from the HVIDB database^51^.

### Protein evolutionary rate analysis

To study the relative evolutionary rate of protein coding sequences and different regions within them, such as their domains or interfaces, we used two separate approaches: (1) dN/dS calculations (the ratio between the number of nonsynonymous and synonymous substitutions) as well as (2) an approach similar to that used by us and others^41,42^ in studies that compare the evolutionary rates between proteins, domains, regulatory and structural regions based on Rate4Site^24^.

In both cases, we used the human protein as a reference and a set of representative one-to-one orthologs, to obtain amino acid substitutions across the vertebrate clade. Orthology relationship and orthologous protein sequences were collected from the annotated genomes of eleven selected vertebrates with relatively high N50 value: Chimpanzee, Crab-eating macaque, Mouse Lemur, Mouse, Rat, Alpine marmot, Dog, Pig, Elephant, Chicken and Abingdon Island giant tortoise using ENSEMBL BioMart (version 102)^61^. The subsequent evolutionary analysis we performed only the genes that have annotated one-to-one orthologs in all 12 species and, in each case, took the longest protein per gene for each species (n=7,445).

For dN/dS calculations, alignments were created for the ortholog protein sequences using PRANK^62^, with masking of unreliably aligned positions in the MSA using GUIDANCE (version 2.02)^63^ and default parameters of 100 bootstrap iterations, sequence default cutoffs of <60% and columns cutoffs of <93% (with the parameters: - -program GUIDANCE2 - - seqType nuc - -msaProgram PRANK - -MSA Param “\-F \-codon”). A phylogenetic tree was built for each MSA using PhyML^64^ (version 3.1). To compute dN/dS values, we used Selecton (version 2.4)^23^ that performs accurate Bayesian rate estimations with prior Bayesian distribution of beta+w. We obtained dN/dS value values for each residue in 7,435 one-to-one orthologs in all 12 species (10 genes were dropped during the run due to having too long sequence).

For Rate4site calculations, alignments were created for the ortholog protein sequences using MAFFT^65^ (version 7.450) with parameters: --anysymbol --maxiterate 1000 – localpair. The resulting multi-sequence alignments (MSAs) were concatenated to form a single MSA, composed of all 7,445 protein MSA files. The MSA was transformed into PHYLIP format using MUSCLE^66^ (version 3.8.31). A phylogenetic tree was built using PhyML^64^ (version 3.1), using the Maximum Likelihood approach with the concatenated alignment of the above-mentioned set of 7,455 proteins, the LG matrix, 4 different substitution rate categories and optimization of tree topology, branch length and rate parameters with the SPR method (running PhyML with the parameters: -d aa -b 0 -o tlr -s SPR). The resulting tree recapitulated the known topology for these 12 vertebrates (**Supp Fig 2**).

To compute the evolutionary rates for each residue in the concatenated MSA, we used Rate4Site^24^ (version 3.0) that performs accurate Bayesian rate estimations, and applied it using the JTT matrix with 8 categories for Bayesian rates and with no branch lengths optimization (with the parameters: -ib -k 8 mj -bn). This yields a residue-specific rate that indicates how fast this residue has evolved across the given clade relatively to the average rate across all resides within the concatenated MSA. The average per protein rates, or per segment within proteins (e.g., specific domains, all surface residues, etc.) were calculated as the mean value of all residues within this protein (or segment).

A correlation between dN/dS values and mean evolutionary rates computed using Rate4site, shows high and significant agreement between the two measures (R =0.901, **Supp Fig 3**).

### Analysis of domains, core, surface and interface residues

We used domain annotations as defined by Pfam-A^25^ to determine regions within proteins that belong to different domains. To compare between the evolutionary rates of different domains – specifically, PRDs *versus* non-PRDs, we took the mean rate of all the residues within these domains. In several cases, such as when comparing PRD domains by types between those that are known to interact with viral proteins and those that are not, we also compared the average rates of any given type of amino acid between the viral-binding PRD and the non-viral-binding PRDs, or included the entire set of residues within the compared distribution.

To define residues as belonging to surface or to core in domains with solved structures (or with AlphaFold-predicted structures^35^), we used the CSU software^67^ to calculate the solvent accessible surface area (ASA). ASA was defined as the ratio of the solvent accessible surface for a given residue within the structured protein versus in the free-state of that residue. Residues were classified as core for ASA ≤ 0.15, and as surface for ASA > 0.15.

To define residues as belonging to the interface between a PRD and its motif (i.e., the residues in the binding pocket of the PRD) we used contacts identified between residues in the PRD and residues in the motif based on the CSU analysis^67^ of the domain-motif PDB structure. The CSU considers atom types, their distance and the interacting surface area for the definition of potentially interacting residues. Those PRD residues found to be interacting with motif residues were subsequently defined as interface residues.

Since the evolutionary rates were computed based on ENSEMBL sequences, while the structural analysis (such as defining core, surface and interface) was often based on PDB structures, we aligned the ENSEMBL-based and the PDB-based sequences using BLASTP (version 2.10.0)^68^ to match the two sequences.

For proteins with no solved structures, we used PredictProtein^69^ (version 1.0.42) software that integrates feature prediction for solvent accessibility (using RePROF). We used PredictProtein definitions for surface and core residues (a 0.16 score as the threshold). To identify interface residues (those that reside in the binding pocket of the PRD and interact with the motif) we used domains with solved structures that include the motif segments, where we identified the interface residues using the CSU method as described above. We then evaluated the sequence similarity of the PRDs with solved structures to homologous PRDs with no solved structure (that belong to the same type of PRD but are within a protein that its structure was not solved) using BLASTP. We kept those PRDs with a similarity of 60% or above in the interface region, and defined the residues that aligned to the interface residues in the solved PRD structures as interface residues in PRDs with no solved structure.

### MSA analysis of SARS-Cov-2 Spike glycoprotein

MSA based on Spike glycoprotein was created using the same approach described above. Sequences were collected from Uniprot of 10 selected Coronaviruses derived from the following accession numbers: SARS-CoV-2: P0DTC2, RaTG13: A0A6B9WHD3, HKU3: A0A7G6UAJ9, WIV1: AGZ48828, RsSHC014:AGZ48806, WIV16: ALK02457, Pangolin-CoV: A0A6M3G9R1, MERS: R9UQ53: SARS-CoV-1 - P59594, RmYNO2: A0A7S8R8N8.

### Within-human protein-protein interaction analysis

We obtained the curated list of physical interactions between human proteins from the STRING database^70^ (version 11.0). For each gene, we used the interactions for its longest protein isoform, and filtered all other isoforms.

### Inference of domain-motif interactions in the set of known protein-protein interactions

For each protein with an annotated PRD (as determined by Pfam-A), we estimated the number of PPIs that are mediated through domain – motif interactions, from the entire set of known PPIs, downloaded from the STRING database. For this, we searched for motifs that are known to be able to specifically bind the PRD in the set of interacting human proteins (e.g., for a protein with an SH3 domain, we searched in its interacting partners whether they include a matching SH3-binding motif). The list of motifs that are known to bind PRDs and their matching PRD was downloaded from the ELM database^50^ (downloaded on 22/03/2021). In each protein that is known to physically interact with a PRD-containing protein, we searched for matching motifs segments. This was done by using the regular expression of these motifs, as defined by the ELM database. We further tested whether the motif matching segments are embedded within a disordered region, by using IUPred^58^ (version 2A, with the ‘long’ option), as explained above. A schematic explanation for this analysis can be found in **Supp Fig 1**. This analysis was used to test how many interacting partners interact with a PRD protein through domain-motif interactions, to compare whether those PRDs that are also bound by viral motifs differ in their connectivity with respect to other PRDs that are not known to be bound by viruses (**Fig 3B**).

### Gene essentiality analysis

We used a dataset of 1,093 essential genes, assembled by Bartha *et al*.^31^. In this work genes were defined as essential using several metrices and included data from *in vitro* and *in vivo* studies. We consider a gene to be essential if it was found to be essential in at least one of the three main screens used in this work.

### Gene expression analysis

We downloaded gene expression data from the human protein atlas^36^. Gene expression level was gathered from 37 tissues of human adult. We calculated the mean expression per gene across tissues (using TPMs – transcript per million measure). In **Supp Fig 9**, we repeated the same analysis as in **Fig 3C**, but this time included only genes that their mean TPM is above 5 or calculated the median value instead the mean value. For the analysis shown in **Fig 4F**, we also computed the variance and median expression across those tissues. Finally, in **Supp Fig 12**, we also show the difference between the gene’s maximal level of expression across tissues and its mean or median expression across these tissues.

### Functional enrichment analysis

We used two different programs to test for functional enrichment of genes of interest: To study the functional enrichment of genes with signatures of positive selection (against the background of the entire human proteome) we used g: Profiler (**Supp Table 4**). To study the functional enrichment of genes in contrast to a chosen background set of genes, we used GOrilla. We used the latter to study enrichment of (1) viral-binding PRDs *versus* non-viral-binding PRD proteins, and (2) viral-binding PRDs *versus* viral-binding non-PRD proteins (**Fig 3E-F**). For the complete list of all enriched terms in these analyses – see **Supp Tables 2-3**.

### Analysis of positively selected genes and residues

To study occurrence of genes with signatures of positive selection in our dataset, we used a previous analysis that identified 331 protein coding genes as high-confidence targets of positive selection in the primate clade, out of 11,170 human genes that had one-to-one orthologs in nine primates^34^. Within these proteins, we used a set of 934 residues that have likely evolved under positive selection.

We studied positively selected residue location in various structural regions (core, surface and interface) of PRD and non-PRD proteins, by overlaying these residues from proteins with solved structures, or with predicted structures from AlphaFold, with their structural analysis, obtained from the CSU software, as explained above. The number of contacts formed between residues within each protein (**intra-contacts**) and between different interacting proteins (**inter-contacts**) was determined using the CSU software.

### Visualizing 3D protein structures

We used the PyMol software^71^ (version 2.0) to visualize and display colored 3D structures shown in **Figs 1B, 2** and **3C**.

### Statistical Analyses

Statistical analyses (Mann-Whitney test, Fisher’s exact test, Spearman’s rank-order correlation and FDR correction based on Benjamini-Hochberg procedure^72^) were performed using the scipy package in Python (version 3.9). Data in boxplots represent the median, first quartile and third quartile with lines extending to the furthest value within 1.5 of the interquartile range. Violin plots show the kernel probability density of the data. Plots were created using matplotlib, seaborn and Plotly.

## Data and script availability

All datasets used in this work (host-virus PPIs, within-human PPIs, scans for positive selection, gene essentiality, gene expression data, orthology relationship, PDB structures, alphaFold predicted structures, PFAM domain, ELM regular expression) are described in the text, including version and time of download, if applicable. The data from these sources were analyzed by processed using Python scripts (version 3.9). Tables containing the metadata, relevant for the analysis, are provided in the supplemental that accompany this manuscript in Excel format (**Supp Table 1-4**). We believe that all data have been described; however, should any additional clarification be needed, the authors will strive to make it available upon request. Please address any additional question to T. Hagai and G. Shuler.

## Code availability

List of command lines for various stages of the protein evolutionary rate analysis, dataset assembly, supplementary tables and protein classification are available on GitHub: https://github.com/galshuler/Evolution_of_Host-Virus_protein_protein_interactions.

## Acknowledgements

We thank Evgeny Fraimovitch, Sivan Friedman-Nakar, Leonid Gitlin, Micahel Lässig, Orly Laufman, Pablo Murcia, Tina Perica and Adi Stern for helpful discussions during the analysis and for comments on the manuscript. This research was supported by the Israel Science Foundation (ISF, grant No. 435/20), by the United States - Israel Binational Science Foundation (BSF, grant No 2019037) and the QBI/UCSF-Tel Aviv University joint Initiative in Computational Biology and Drug Discovery.

