## Supporting Figures for "Rapidly evolving viral motifs target biophysically constrained binding pockets of host proteins"

### Supporting information:

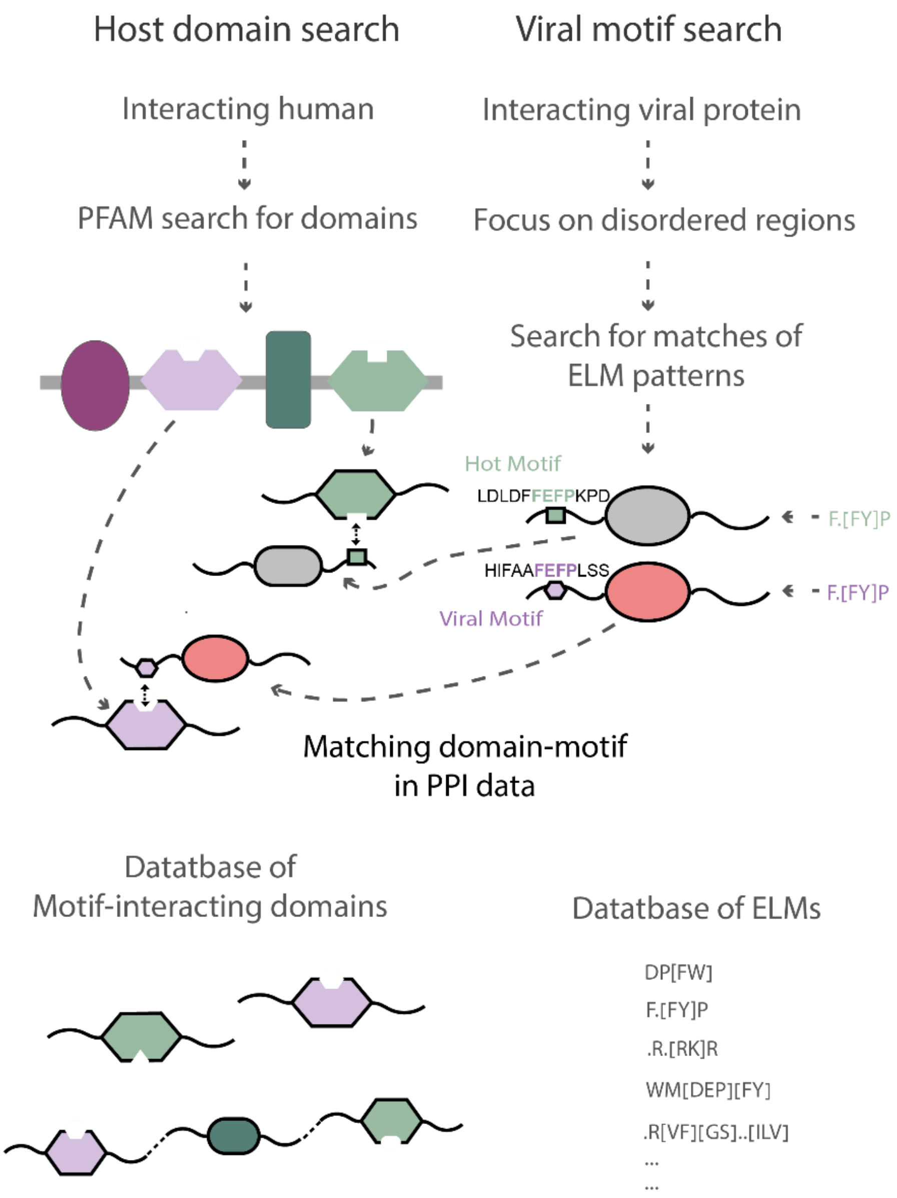


**Supporting Figure 1. A schematic of the methodology to infer domain-motif interactions**. For each human protein with an annotated PRD (as determined by Pfam-A), we searched for the PPIs that are potentially mediated through domain – motif interactions, from the entire set of known PPIs, downloaded from the HVIDB database. For this, we searched for motifs that are known to be able to specifically bind the PRD in the set of interacting human proteins (e.g., for a human protein with an SH3 domain, we searched in its interacting viral partners whether they include a matching SH3-binding motif). The list of motifs that are known to bind PRDs and their matching PRD was downloaded from the ELM database (downloaded on 22/03/2021). We further tested whether the motif-matching segments are embedded within a disordered region and excluded matches that are found in ordered regions. The example above describes inference of host domain – viral motif interaction, however the same procedure was also used for within-human PPIs to infer host domain – host motif interactions.

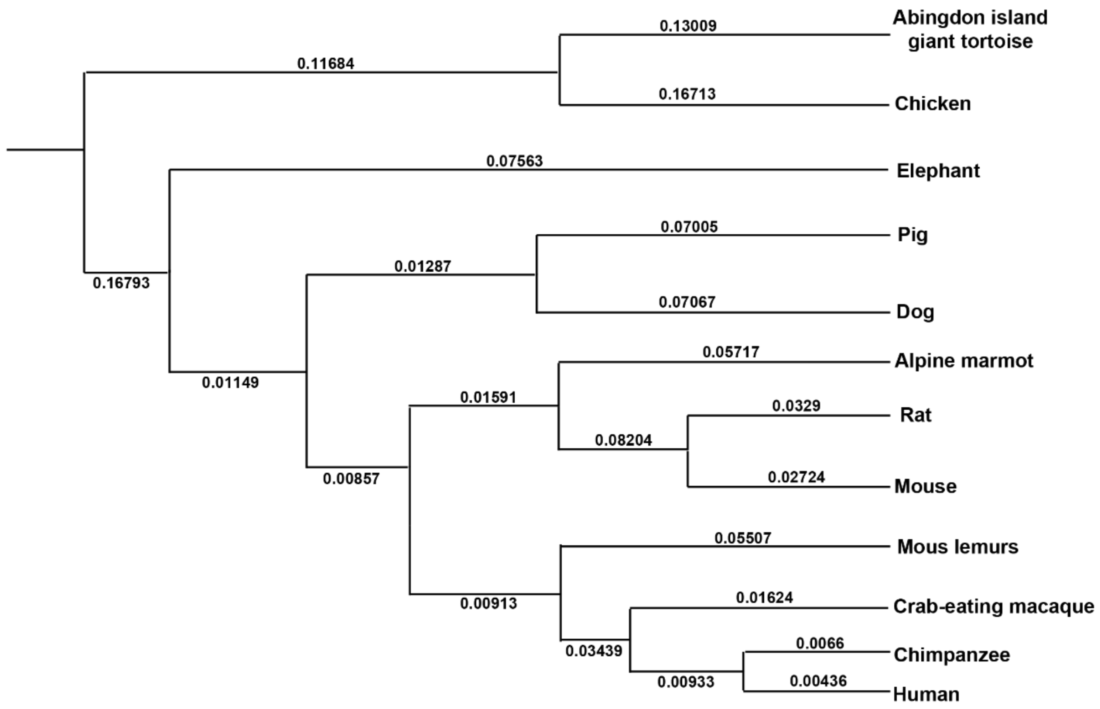


**Supporting Figure 2. A schematic of rectangular phylogenetic tree of our one-to-one orthologs**. A phylogenetic tree with branch length optimization (indicated at the center of each branch; branch length was ignore) was built using PhyML (version 3.1) based on protein sequences that were collected from the annotated genomes (ENSEMBL, version 102) of twelve selected vertebrates with relatively high N50 value: Human,Chimpanzee, Crab-eating macaque, Mouse Lemur, Mouse ,Rat, Alpine marmot, Dog, Pig, Elephant, Chicken and Abingdon island giant tortoise, using the Maximum Likelihood approach with the concatenated alignment of the above-mentioned set of 7,455 proteins. The tree recapitulated the known phylogenetic relationship for these species.

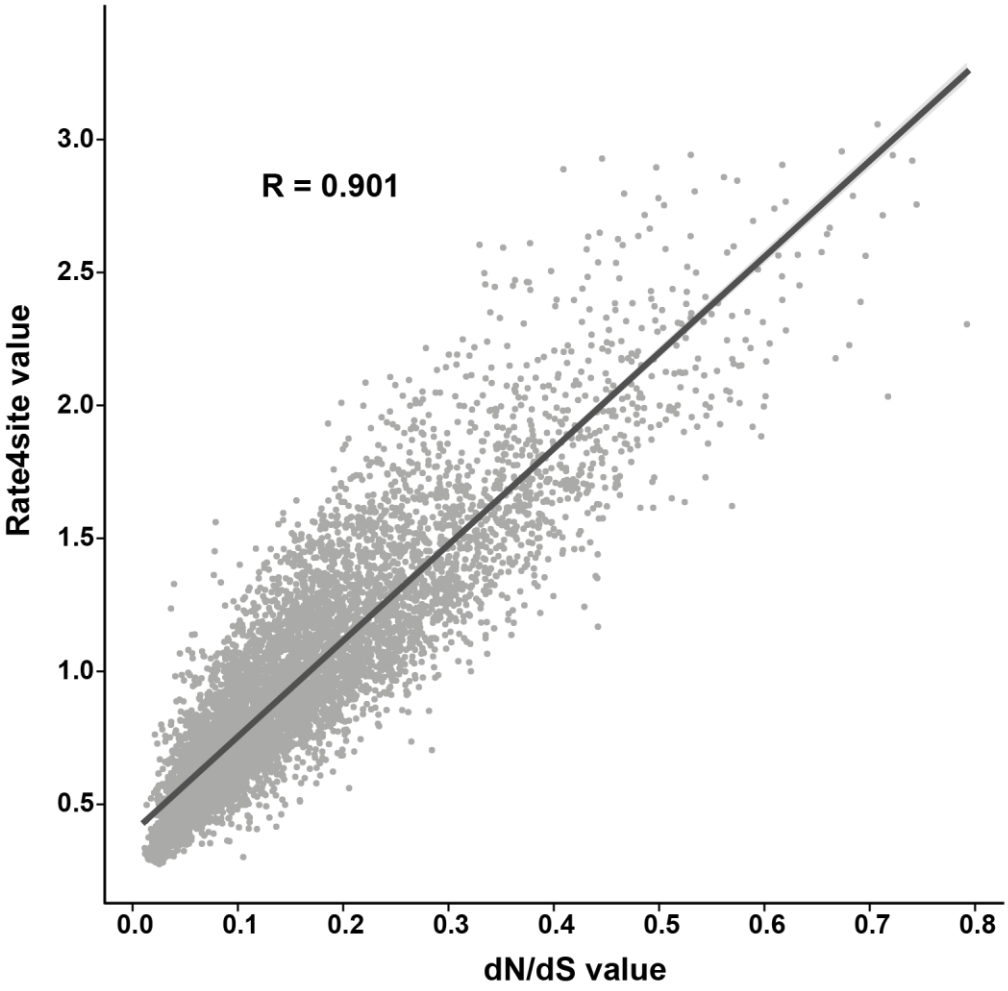


**Supporting Figure 3. Correlation between dN/dS and Rate4site measures.** Scatter plot of evolutionary rates in human proteins, computed with additional 11 one-to-one orthologs from non-human vertebrates (n = 7,435). The Y-axis represents the average evolutionary rate across all residues of the protein, calculated with Rate4site, and the X-axis represents the dN/dS values of the proteins, calculated with Selecton. A significant correlation is observed (R = 0.901, P-value$<10^{-308}$, Spearman's rank correlation test).

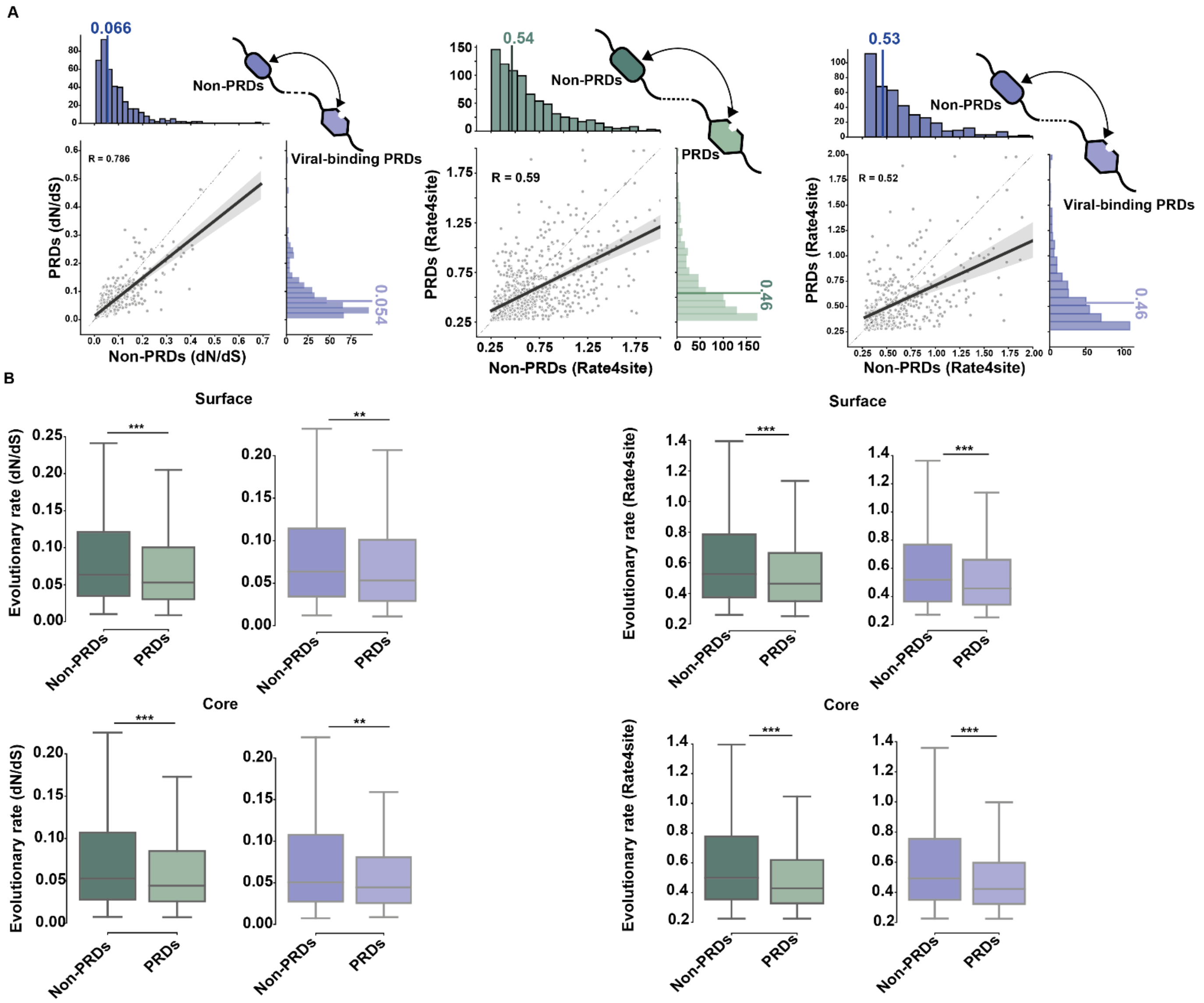


**Supporting Figure 4. Comparison of evolutionary rates between domains and between structural regions in the same protein. (A)** Paired distributions of evolutionary rates of domains within proteins, where the Y-axis represents the average value of the PRDs (green) or PRDs known to interact with viruses (purple) and the X-axis represents the average value of the non-PRDs from the same protein (n = 410 proteins). PRDs have significantly lower average rate in comparison to non-PRDs (left – dN/dS approach, middle and right – Rate4site approach, Mann-Whitney test). **(B)** Distribution of evolutionary rates of domains within proteins, partitioned into core and surface residues. Boxplots represents the average value of the PRDs (green) or PRDs known to interact with viruses (purple) compared to non-PRDs. The differences between the distributions were compared using Mann-Whitney test. (left – dN/dS approach, right – Rate4site approach, *** - P < 0.001, ** -P < 0.01, * P < 0.05).

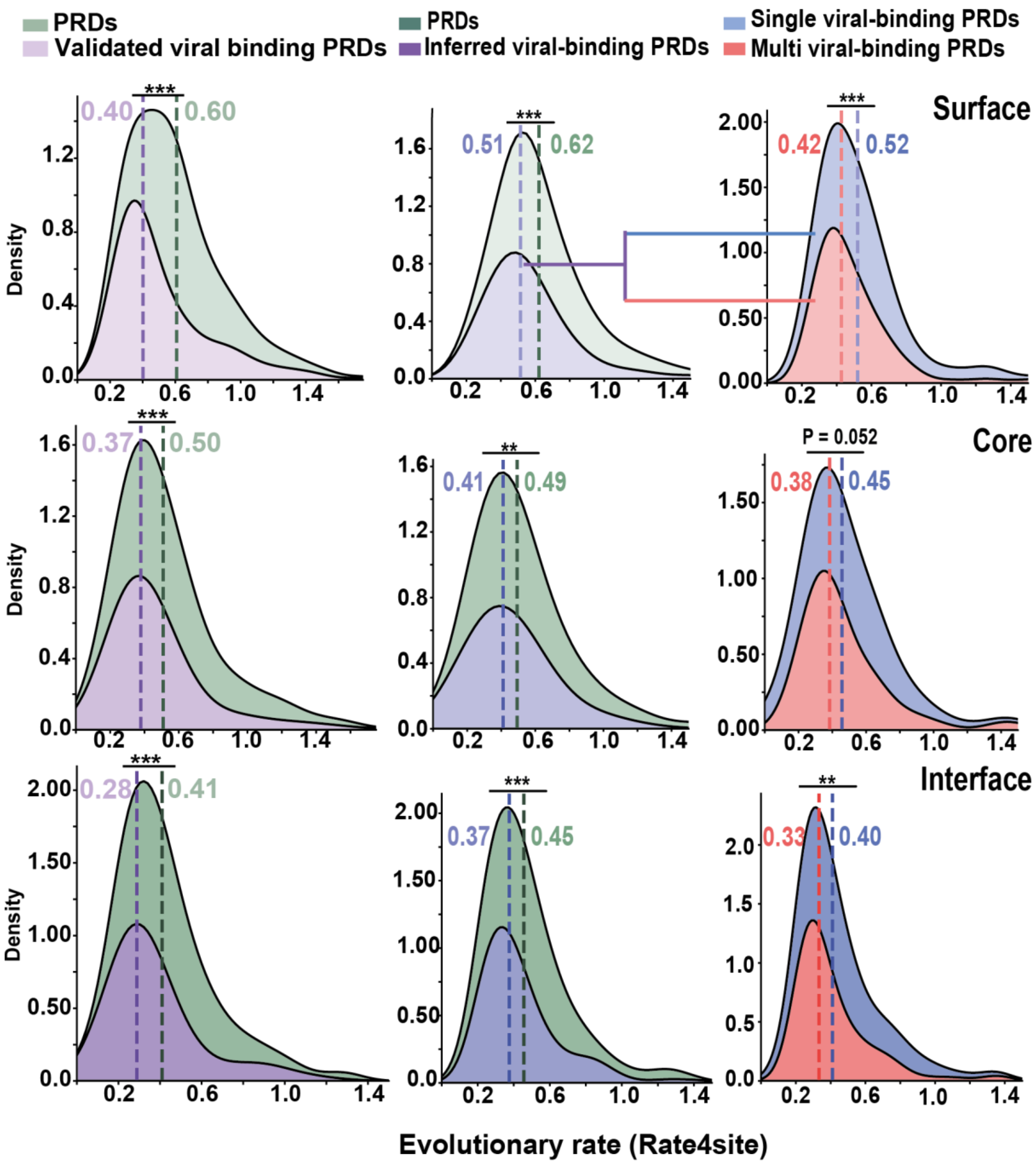


**Supporting Figure 5. Distribution of evolutionary rates of matched amino acids at the core, surface and interface of PRDs using the Rate4site approach**. Kernel density plots showing the distribution of evolutionary rates of amino acids at the core, surface and interface of PRDs (left - experimentally validated viral-binding PRD, middle and right - inferred viral-binding PRDs). PRDs are split into those that are not known to be bound by viral motifs (left and middle, in green) those that are known to be bound viruses through domain-motif interaction (left and middle, in purple) and those by those that are known to be bound by a single (right, in blue) and multiple viral motifs (right, in red). Dashed lines and values represent the medians of the distribution. The differences between the distributions were compared using Mann-Whitney test. (*** - P < 0.001, ** -P < 0.01, * P < 0.05).

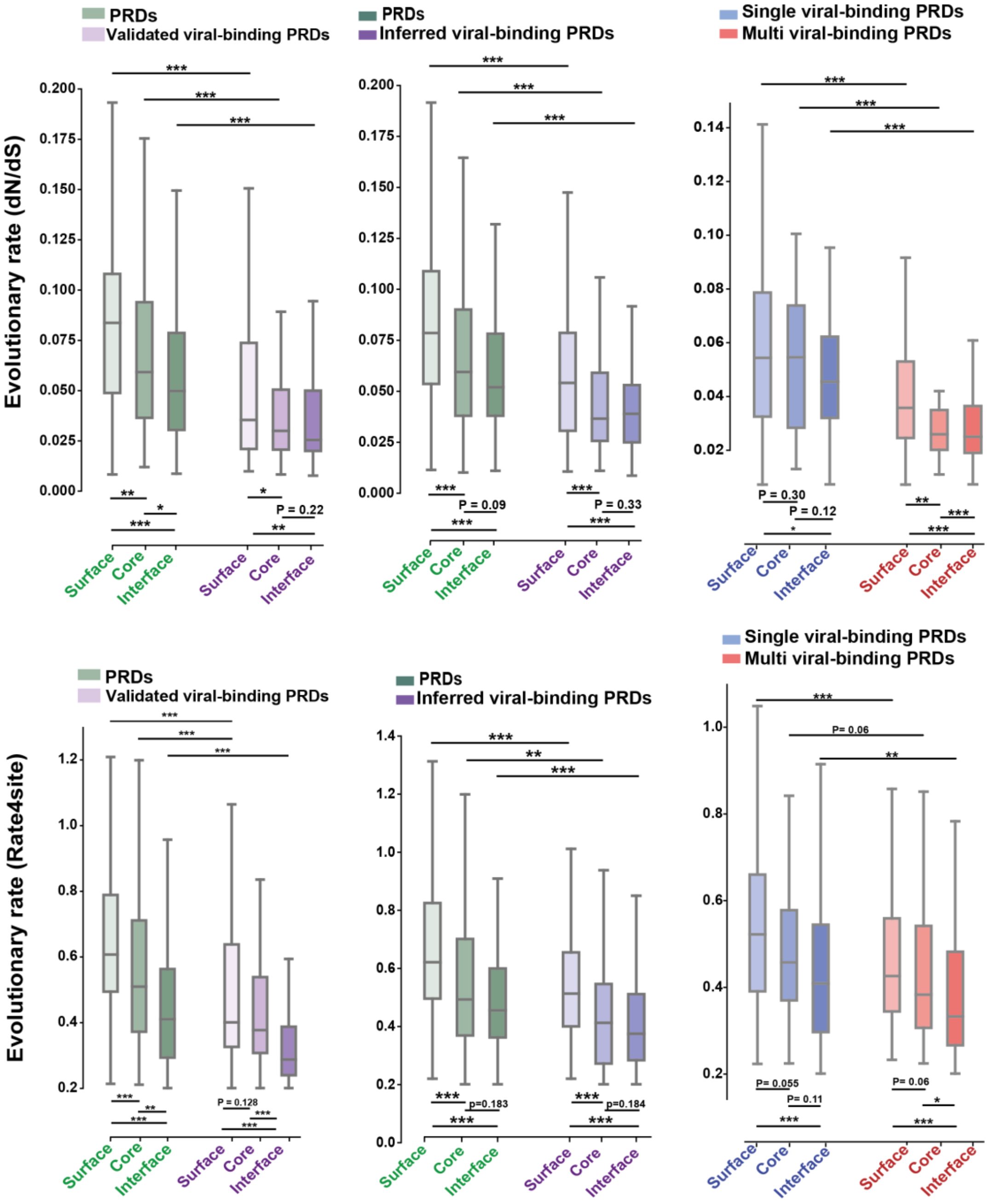


**Supporting Figure 6. Distribution of evolutionary rates of matched amino acids at the core, surface and interface of PRDs.** As shown in Fig 1F and Supporting Figure 5, PRDs are split into those that are known or not known to be bound by viral motifs (left and middle, in purple and green respectively) and those that are known to be bound by a single and multiple viral motif (right, in blue and red respectively). The differences between and within the distributions were compared using Mann-Whitney test and corrected by FDR. (top – dN/dS approach, bottom – Rate4site approach, *** - P < 0.001, ** -P < 0.01, * P < 0.05).

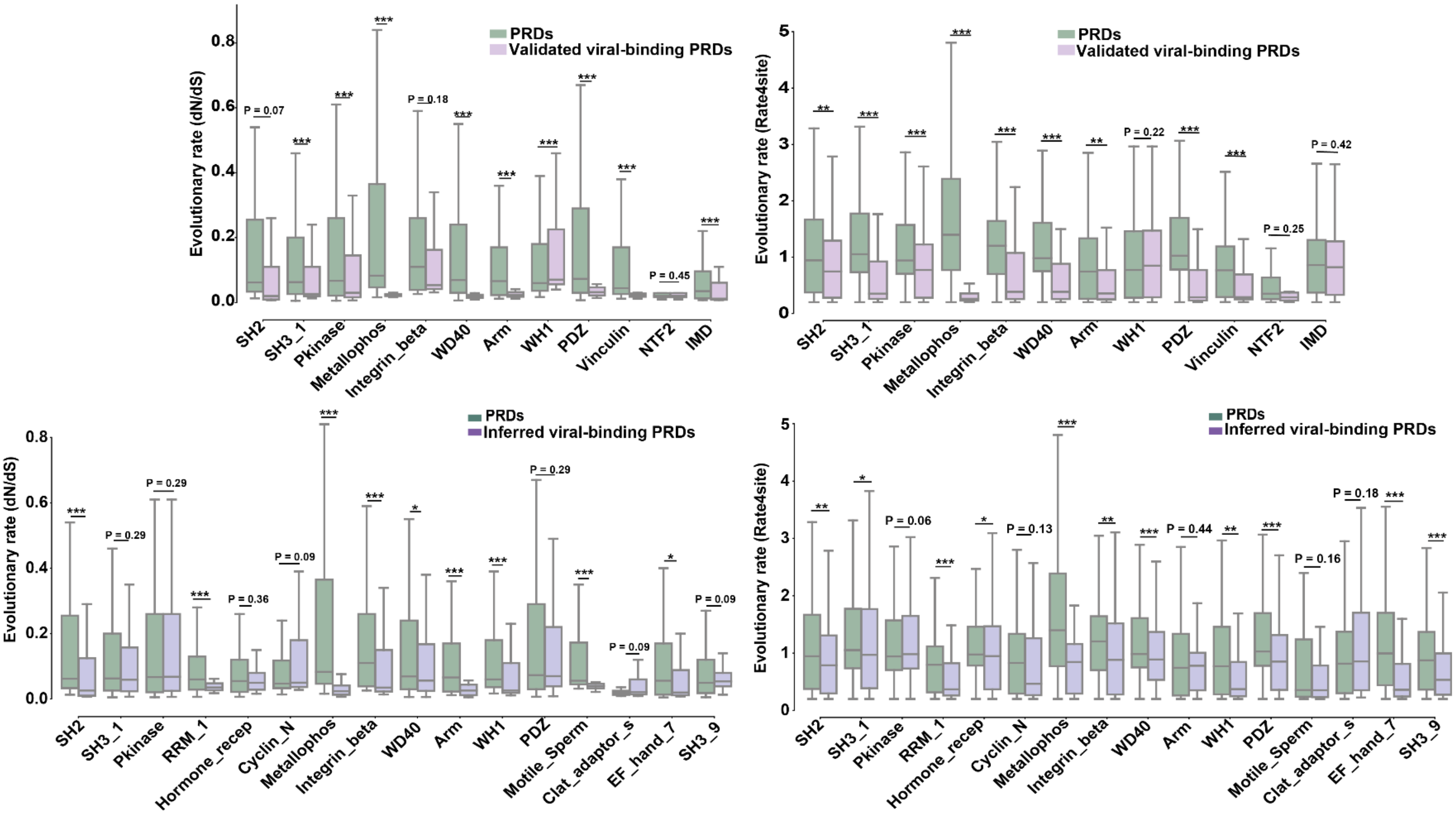


**Supporting Figure 7. Distribution of evolutionary rates of all amino acids within PRDs, partitioned by a PRD type.** PRDs are partitioned by the PRD type. In each PRD type, the PRDs are split into those that are known or not known to be bound by viral motifs (top and bottom, in purple and green, respectively, including both inferred and experimentally validated sets). The differences between the distributions were compared using Mann-Whitney test and corrected by FDR. (left – dN/dS approach, right – Rate4site approach, *** - $P < 0.001$, ** -$P < 0.01$, * $P < 0.05$).

A

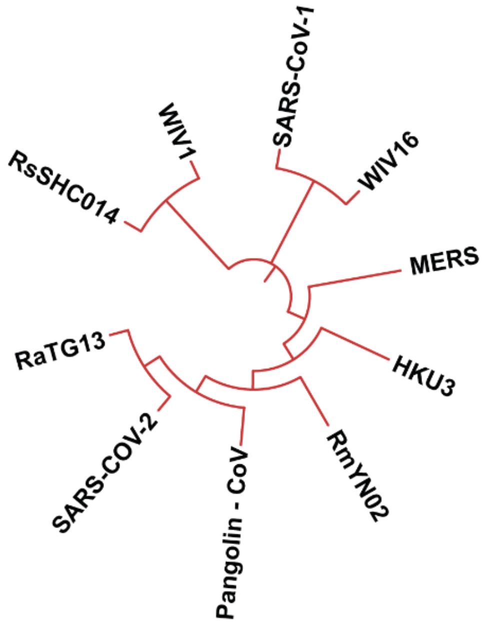


B

| | 296 | | | | | | 302 | 315 | | | | | | 321 | 345 | | | | | | | | | 354 |
|---|---|---|---|---|---|---|---|---|---|---|---|---|---|---|---|---|---|---|---|---|---|---|---|---|
| Human | Q | Y | S | T | N | W | S | W | T | P | G | E | D | S | I | S | K | E | T | K | K | K | Y | Y |
| Chimpanzee | Q | Y | S | T | N | W | S | W | T | P | G | E | D | S | I | S | K | E | T | K | K | K | Y | Y |
| Mouse | Q | Y | G | T | N | W | S | W | T | P | G | E | D | S | I | S | K | E | T | K | K | K | Y | Y |
| Pig | Q | Y | S | T | N | W | S | W | T | P | G | E | D | S | I | S | K | E | T | K | K | K | Y | Y |
| Horseshoe bat | Q | Y | S | T | N | W | S | W | T | P | G | E | D | S | I | S | K | E | T | K | K | K | Y | Y |
| Megabat | Q | Y | S | I | N | W | S | W | T | P | G | E | D | S | I | S | K | E | T | K | K | K | Y | Y |
| Chicken | Q | Y | S | A | I | W | S | W | T | P | G | E | D | S | I | S | K | E | T | K | K | E | Y | Y |
| Tortoise | Q | Y | S | T | N | W | S | W | T | P | G | E | D | S | I | S | K | E | T | K | R | K | Y | Y |

**Supporting Figure 8. SARS-Cov-2-specific motif binds to the highly conserved binding pocket with Neuropilin Receptor-1 Domain**. **(A)** phylogenetic tree with branch length optimization was built using RaxML[1] (version 8) based on Spike glycoprotein amino acid sequences that were collected from Uniprot of 10 selected corona viruses derived from the following accession numbers: SARS-CoV-2: P0DTC2, RaTG13: A0A6B9WHD3, HKU3: A0A7G6UAJ9, WIV1: AGZ48828, RsSHC014:AGZ48806, WIV16: ALK02457, Pangolin-CoV: A0A6M3G9R1, MERS: R9UQ53: SARS-CoV-1 - P59594, RmYNO2: A0A7S8R8N8, using the Maximum Likelihood approach. **(B)** MSA of residues in the binding pocket of NRP1 across vertebrates, based on an MSA of 8 orthologs from ENSEMBL annotated genomes used in this study and two bat species (Human, Chimpanzee, Mouse, Pig, Horseshoe bat, Megabat, Chicken and Tortoise). In purple – residues in NRP1 domain that interact with the SARS-Cov-2 Motif. See comparison with respective distribution in Fig 2.

1 Alexey M. Kozlov, Diego Darriba, Tomáš Flouri, Benoit Morel, and Alexandros Stamatakis (2019), RAxML-NG: A fast, scalable, and user-friendly tool for maximum likelihood phylogenetic inference

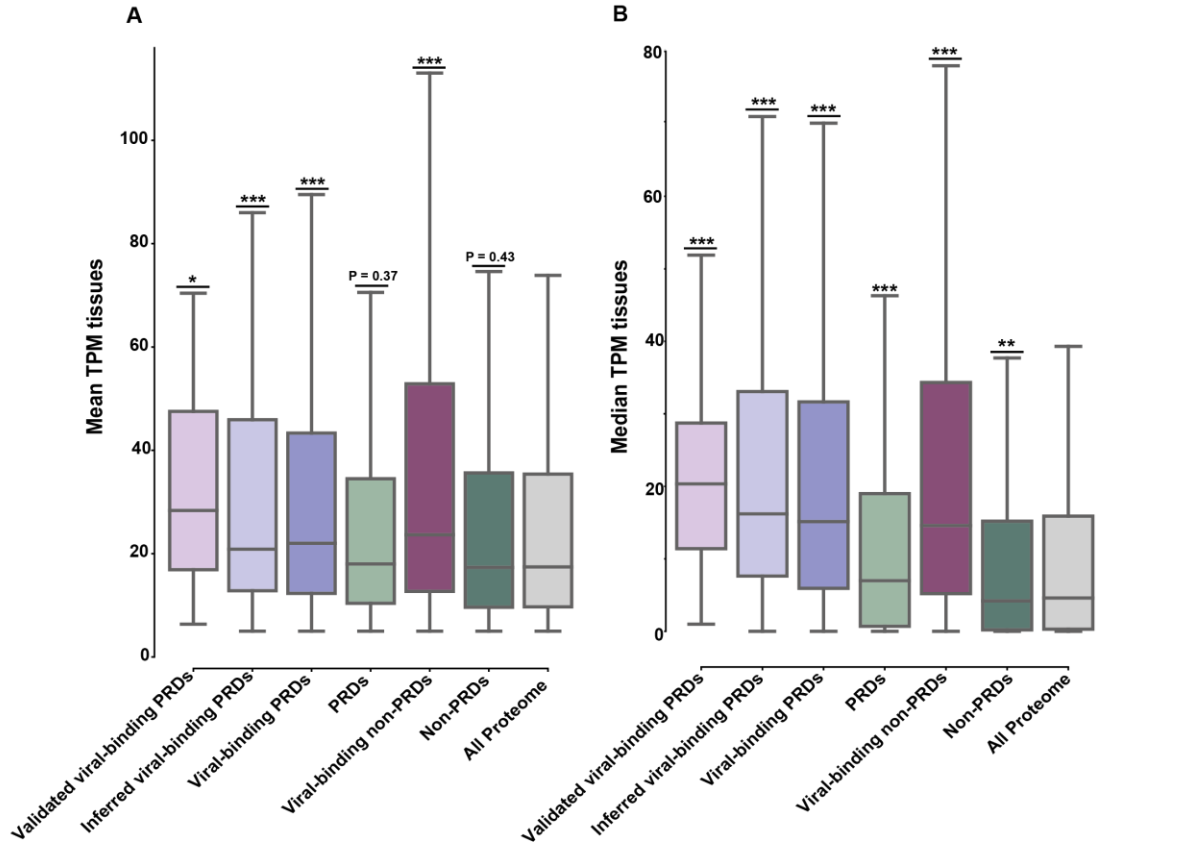


**Supporting Figure 9. Distributions of average gene expression levels across healthy adult human tissues. (A)** As in Fig 3C, but including only genes that their mean TPM is above 5 (n = 11,895). Differences between the distributions were compared using Mann-Whitney test and corrected by FDR. **(B)** Distributions of median gene expression levels across healthy adult human tissues (n = 19,670, in here we include all genes and show median distributions to compare with the average that is shown in Fig 3C). Differences between the distributions were compared using Mann-Whitney test and corrected by FDR. See comparison with respective distribution of mean TPM in Fig 3C. (*** - $P < 0.001$, ** -$P < 0.01$, * $P < 0.05$).

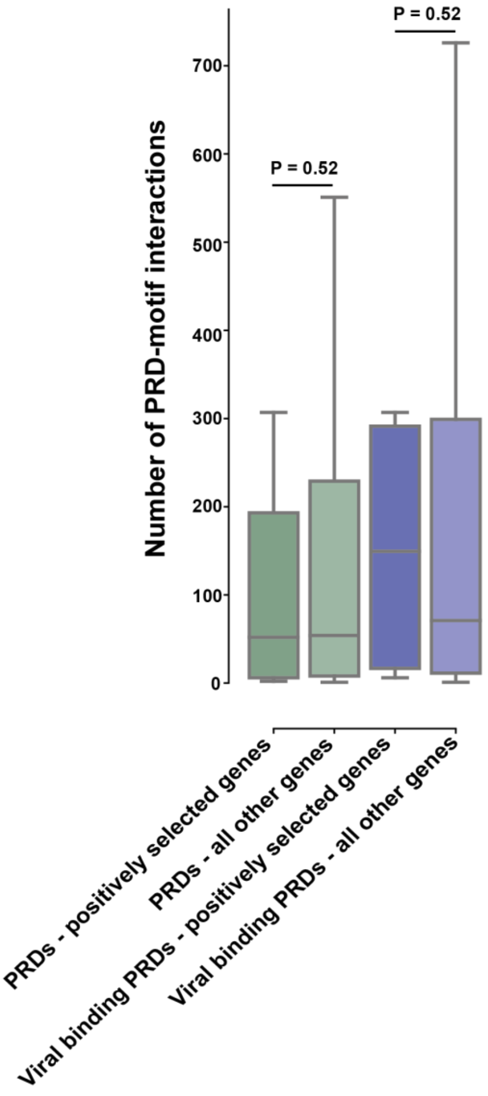


**Supporting Figure 10. Positively selected PRD proteins that bind viruses are not depleted of interactions with other human proteins**. A comparison of "binding pocket usage" between all PRD proteins with and without signatures of positive selection (green) and the subset of PRDs known to bind viral (purple). The differences between the distributions were compared using Mann-Whitney test and corrected by FDR. (*** - P < 0.001, ** -P < 0.01, * P < 0.05). See comparison with respective distribution of the number of PPI within human proteome in Fig 4D.

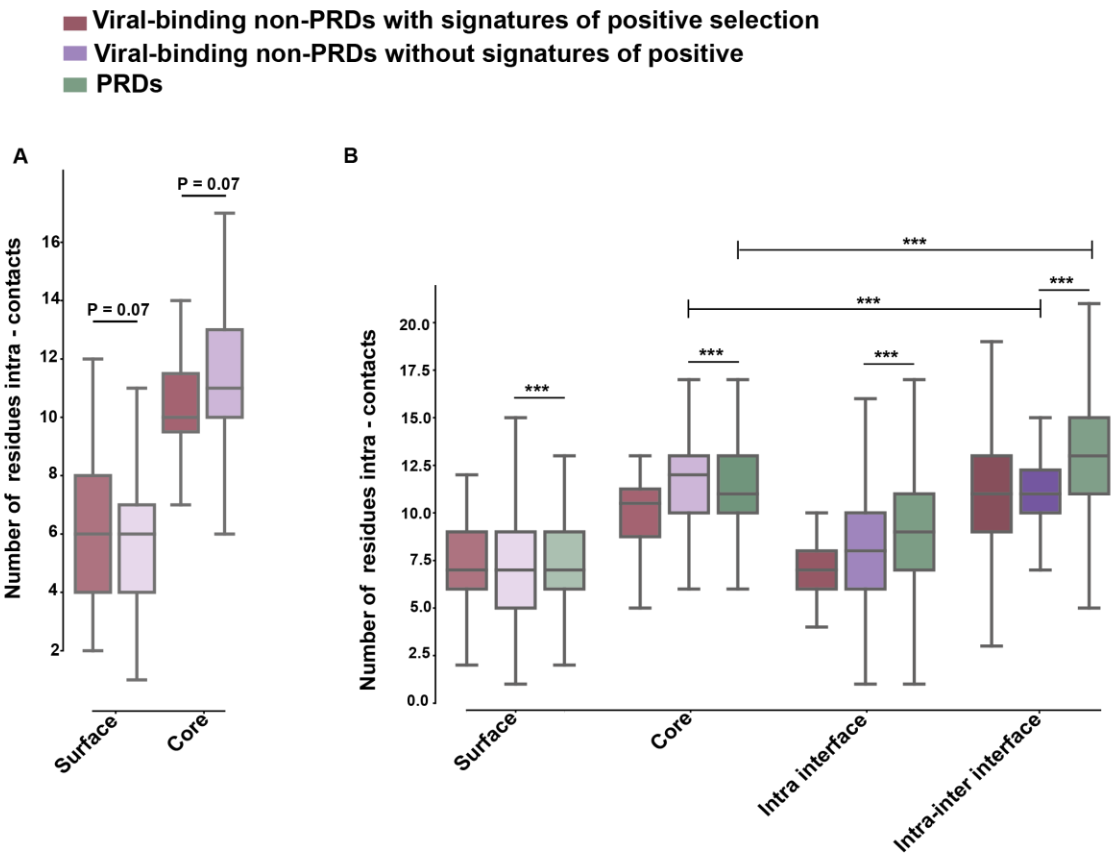


**Supporting Figure 11. Distributions of interactions formed between residues within the same protein (intra-contacts) in different sets of proteins. (A)** Number of intra-contacts in surface and core regions, based on structures predicted by AlphaFold (44 proteins), between viral-binding non-PRDs, with and without signatures of positive selection (dark red gradient and purples gradient, respectively). The differences between the distributions were compared using Mann-Whitney test and corrected by FDR. **(B)** Number of intra-contacts in surface, core and interface. In interface regions we compute both “intra contacts” – those interactions formed only within the same domain, and “intra-inter contacts”, which is the combination of contacts that residues at the interface form within the same domain they belong to as well as with residues from the protein they interact with. These different regions are compared between viral-binding non-PRDs and PRD, with and without signatures of positive selection (red, purple and green, respectively). The differences between the distributions were compared using Mann-Whitney test and corrected by FDR (*** - P < 0.001, ** -P < 0.01, * P < 0.05). See comparison with respective distribution of viral-binding non-PRDs in Fig 4E.

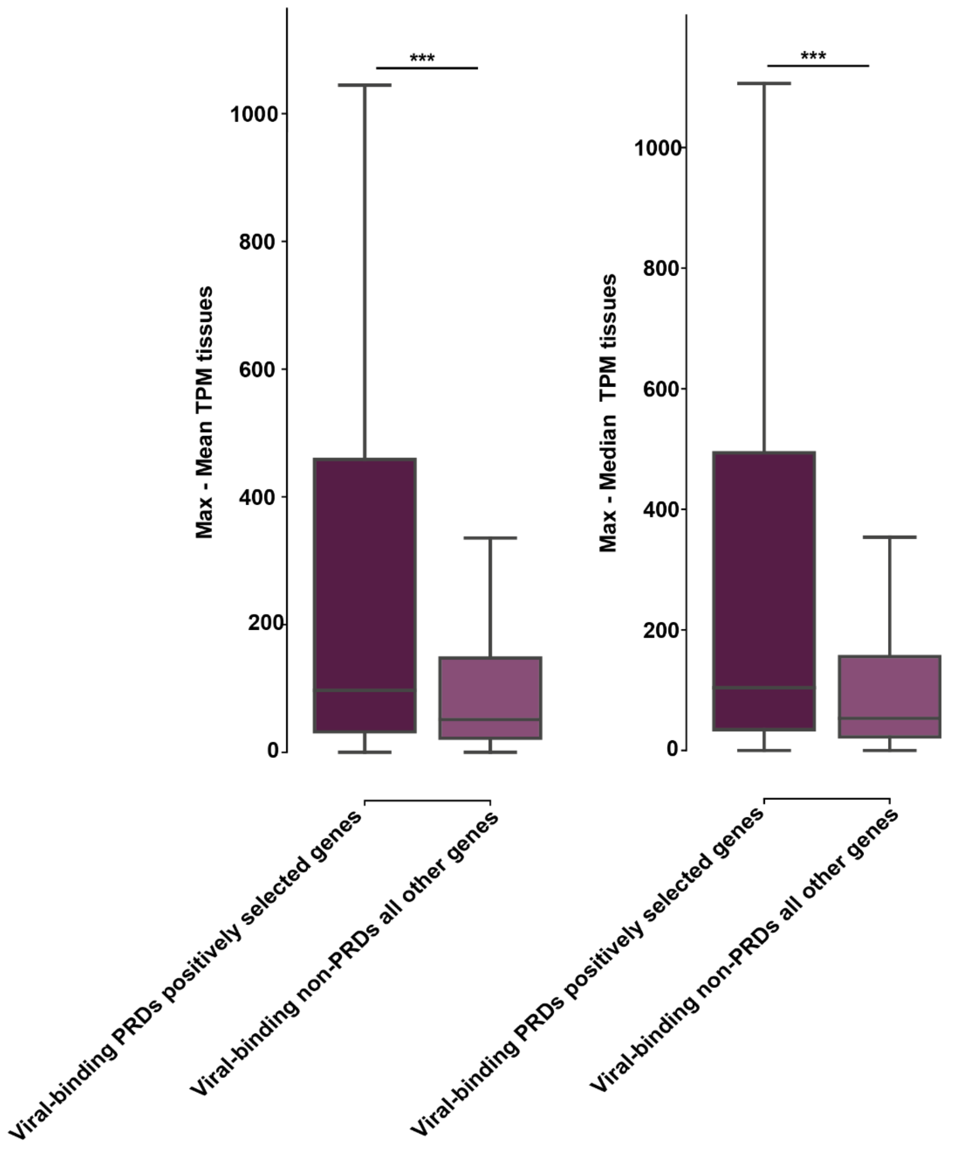


**Supporting Figure 12 Comparisons of measures of cross-tissue gene expression between viral binding non-PRDs with and without signatures of positive selection**. Left - Max value minus mean value of TPMs, right – Max value minus median TPMs (where Max is the maximum TPM value found across the tissues for each gene). The differences between the distributions were compared using Mann-Whitney test and corrected by FDR. Positively selected genes show higher Max-Median and Max-Mean in comparison with other genes. (*** - P < 0.001, ** -P < 0.01, * P < 0.05). In all cases, positively selected genes are significantly higher than other genes, except when comparing the median distributions (in Fig 4F, middle). This suggests a tissue-specific expression of one or few tissues where the positively selected genes are more highly expressed than the other genes. See comparison with respective distribution of mean, median and variance TPM tissues in Fig 4F.
